# Microfluidic creep experiment for measuring linear viscoelastic mechanical properties of microparticles in a cross-slot extensional flow device

**DOI:** 10.1101/2024.08.07.607090

**Authors:** Sara Ghanbarpour Mamaghani, Joanna B. Dahl

**Affiliations:** Engineering Department, University of Massachusetts Boston, Boston, MA 02025, USA

**Keywords:** Microfluidics, Microrheology, Creep

## Abstract

The micromechanical measurement field has struggled to establish repeatable techniques, likely because the deforming stresses can be complicated and difficult to model. Here we demonstrate experimentally the ability of cross-slot microfluidic device to create a quasi-steady deformation state in agarose hydrogel microparticles to replicate a traditional uniaxial creep test at the microscale and at relatively high throughput. A recent numerical study by Lu et al. [Lu, Guo, Yu, Sui. *J. Fluid Mech.*, 2023, 962, A26] showed that viscoelastic capsules flowing through a cross-slot can achieve a quasi-steady strain near the extensional flow stagnation point that is equal to the equilibrium static strain, thereby implying that continuous operation of a cross-slot can accurately capture capsule elastic mechanical behavior in addition to transient behavior. However, no microfluidic cross-slot studies have reported quasi-steady strains for suspended cells or particles, to our knowledge. By using large dimension cross-slots relative to the microparticle diameter, our cross-slot implementation created an extensional flow region that was large enough for agarose hydrogel microparticles to achieve a strain plateau while dwelling near the stagnation point. This strain plateau will be key for accurately and precisely measuring linear viscoelastic properties of small microscale biological objects. The mechanical test was performed in the linear regime, so an analytical mechanical model derived using the elastic-viscoelastic correspondence principle was proposed to extract linear viscoelastic mechanical properties from observed particle strain histories. Particle image velocimetry measurements of the unperturbed velocity field were used to determine where in the device particles experienced extensional flow and the mechanical model should be applied. The measurement throughput in this work was 1 – 2 particles achieving a quasi-steady strain plateau per second, though measurement yield and throughput can be increased with particle-centering upstream device design features. Finally, we provide recommendations for applying the cross-slot microscale creep experiment to other biomaterials and criteria to identify particles that likely achieved a quasi-steady strain state.

## INTRODUCTION

Many studies have shown the clinical potential of using high-throughput microfluidics to mechanically phenotype or group cells based on their mechanical behavior for disease diagnostics and therapeutics [1,2]. To measure intrinsic cell mechanical properties, a mechanical model is applied so that cell stiffnesses can be inferred from the observed cell strains under a known deforming force, thereby disentangling cell size and deformation. Otherwise, cell deformation depends on its size, with larger cells tending to stretch more than smaller cells [3,4]. In contrast, intrinsic mechanical properties do not depend on cell size and can facilitate more direct biomechanics comparisons between cells. A challenge for accurate mechanical property measurements is achieving a close match between the mechanical model and the experimental conditions. Many previous microfluidic studies that measured cell mechanical properties relied on asymptotic approximations [5] and empirical calibration [6,7] to estimate the forces that deformed the cells. Others applied phenomenological mechanical models [8] to extract mechanical properties. To make further progress in cell mechanophenotyping, further work is needed in the practical implementation of microfluidic devices for more accurate mechanical property measurements.

Cross-slot extensional flow devices are widely used devices that have under-explored capabilities for measuring cellular mechanical properties. This device design stretches cells into a prolate ellipsoid at the stagnation point (zero velocity) of two impinging flows delivered through two opposing perpendicular straight channels. Cross-slots have been used in the soft matter field to study the extensional rheology of polymeric fluids [9–11], single DNA molecule dynamics [12–15], and synthetic vesicles shape dynamics in extensional flow [16–18]. In the area of cell mechanics, the cross-slot has been used to mechanophenotype cell populations found in patient biofluids [19–21] and to determine population-averaged mechanical properties of cultured cells [5,8]. Cell mechanophenotyping based on 2D scatter plots of cell deformation vs. cell projected area has important clinical relevance and is the basis of a recently approved biomedical device for early detection of sepsis [21]. Less progress has been made in single cell mechanical property measurements using the cross-slot device. Population-averaged viscoelastic stiffnesses have been extracted from the averaged strain histories of many cells [5] and from the average maximum strain of many cells at several different flow rates [8]. To our knowledge, the viscoelastic mechanical properties of individual cells measured in a cross-slot have not been reported.

A recent simulation study suggested practical considerations for performing single cell viscoelastic mechanical property measurements in a cross-slot device [22]. The authors studied the deformation of liquid-filled viscoelastic capsules flowing through a cross-slot device at low and high Reynolds numbers. Consistent with other simulations of solid particles in cross-slots [23], Lu et al. found that capsules not aligned to the device’s cross-sectional geometric center had lower maximum deformations in the cross-slot extensional flow region than well-aligned capsules due to experiencing lower fluid stresses. Therefore, techniques that extract mechanical properties from the cell maximum deformation [5,8] could yield inaccurate elastic moduli if cells are too offset from the device center because offset cells will not reach their steady-state deformed shapes. For low Reynolds number flows (Re ≤ 1), the Lu et al. demonstrated that capsules sufficiently centered (offset less than or equal to 1% of the channel width) achieved a quasi-steady deformation state with a strain (Taylor deformation parameter) that was indistinguishable to the steady-state strain for a capsule held in a fixed position at the stagnation point. This suggests that accurate elastic moduli can be measured for sufficiently centered capsules.

The study of Lu et al. implies that particles and cells could be measured for their creep-like response to a suddenly applied viscous extensional forces in a transient cross-slot system—if they are sufficiently aligned to the center streamline so that they achieve the quasi-steady deformation state. Cross-slots have been used to trap vesicles at the extensional flow’s stagnation point long enough for them to achieve a steady-state deformation using manual trapping for minutes [16,17] and up to hours using computer-controlled pneumatic valves [24]. However, trapping reduces the throughput of cross-slot mechanical property measurements. To our knowledge, studies using cross-slots on biological suspended cells without trapping have not demonstrated a quasi-steady shape as the cells dwelled within the extensional flow region at high flow rates and finite Reynolds numbers [19,20] or at lower flow rates and vanishing Reynolds numbers [5,8]. In those works, the size of the device channels was similar to the suspended cells; the ratio of channel width to cell diameter in the studies varied 1.25 – 3. This meant that the size of the extensional flow cross-slot region in the middle of the device (*w* x *w* area, where *w* is channel width) was of similar size to the cell. In these cases, cells did not have time to achieve a quasi-steady strain state as they passed through the relatively small extensional flow area. Though Lu et al. simulated similar conditions with the square channel width to capsule diameter ratio being 2.5 and predicted quasi-steady strain states for capsules offset <1% channel width [22], the reported experimental implementations suggest that practically no cells achieved a quasi-steady strain state when channel dimensions and cell diameters were of similar size.

Experimental demonstrations of a quasi-steady deformation state for well-aligned capsules or particles would be helpful for developing the cross-slot’s biomechanics measurement capabilities. A quasi-steady deformation state is required for accurately measuring the elastic part of an object’s mechanical response. Reliably achieving this state during transient operation of a cross-slot would be a good compromise between mechanical property measurement accuracy and measurement throughput. Furthermore, mechanical models are needed to relate the observed quasi-steady stretched state of spherical objects and the extensional flow fluid forces to the desired mechanical properties.

Here we present a microfluidic cross-slot technique to measure linear viscoelastic mechanical properties of single microparticles in non-contact extensional flow conditions, an implementation that is reminiscent of a traditional uniaxial creep test. Using the strategy of generating a larger extensional flow region to increase particle dwell times, we demonstrate the presence of a quasi-steady deformation state for agarose microparticles (diameters typically 25-50 µm) in a relatively large cross-slot device (cross-section approximately 290 µm wide, 265 µm deep). We catalog the particle trajectory conditions under which agarose particles tend to achieve quasi-steady deformation while dwelling near the stagnation point. Experimental measurements of the unperturbed fluid velocity field provide insight into the cross-slot flow regimes and where the particles start to feel extensional flow. Using the elastic-viscoelastic correspondence principle, we propose an analytical mechanical model for extracting linear viscoelastic mechanical properties of single spherical particles in a cross-slot. We demonstrate our analysis and experimental approach using agarose hydrogel microparticles that are modeled as a Kelvin-Voigt viscoelastic solid and report distributions of viscoelastic material parameters for individual particles. Finally, we provide recommendations for applying the high-throughput cross-slot microscale creep test for other biomaterials.

## METHODS

### Extensional Flow Microfluidic Device

A microfluidic extensional flow device, commonly called a cross-slot, consists of two channels that intersect at 90 degrees (**Figure 1A**). Within the channel intersection region, approximately planar extensional flow field was generated with a stagnation point (zero velocity) at the center (**Figure 1B**). PDMS devices were prepared using standard soft lithography techniques [25]. Sheathing flow streams intersected the central sample inlet channel at 45° angles with the purpose of centering microparticles as close as possible to the center streamline. The channel length from the sheathing and center intersection to the cross-slot edge was more than 10 mm to ensure fluid flow has fully developed by the location of creep measurements. Device masters were made from laminating three layers of 100-µm thick dry film photoresist (Riston GoldMaster GM130 photoresist, DuPont) on stainless steel wafers (Stainless Steel Supply) [26]. Channel cross-section dimensions were approximately 290 µm wide and 265 µm deep as measured from two optical microscopy approaches: from brightfield microscopy images without fluid in the device and from fluorescence microscopy images with a dense suspension of fluorescent tracer particles infused in the device. The same microfluidic device was used for all hydrogel particle creep experiments and particle image velocimetry measurements.

**Figure 1.**
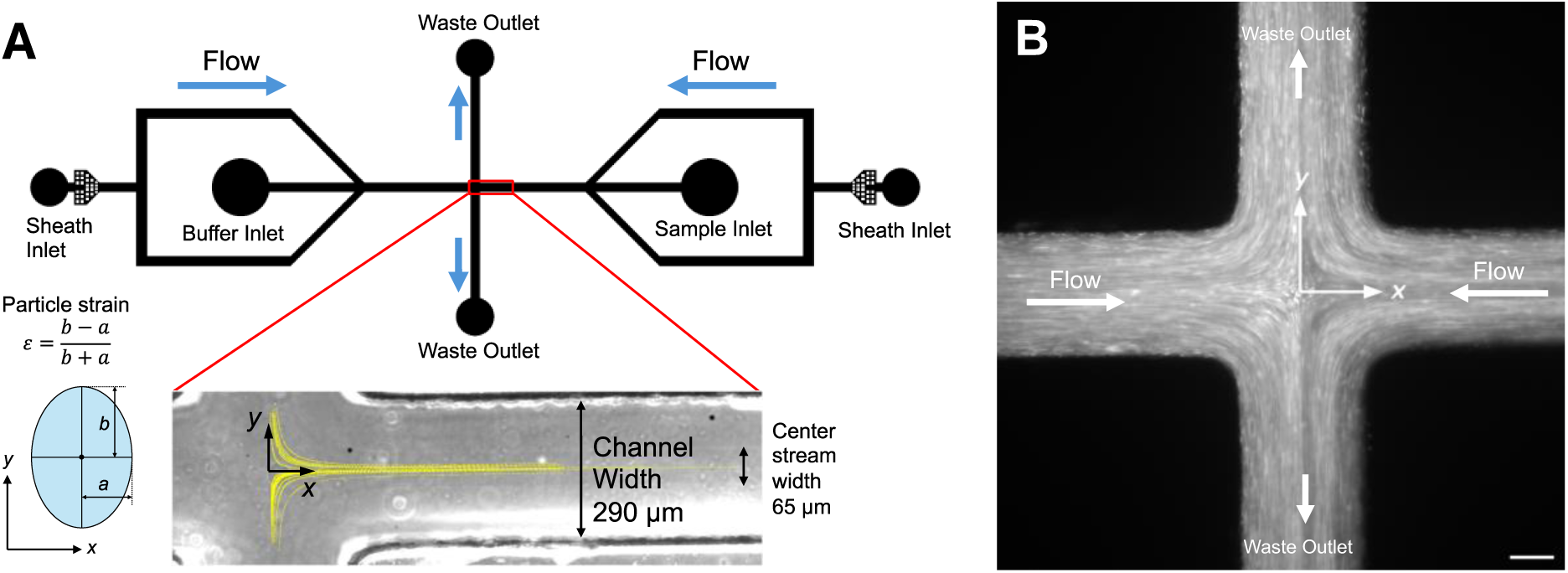
Microfluidic device for microscale creep measurements. **(A)** The cross-slot device with sheathing flow directed particles towards the stagnation point at the center of the extensional flow region. The origin was the stagnation point. The *x* axis was parallel to the inlet flow direction, and the *y* direction was parallel to the outlet flow direction. Overlayed on the image showing the camera field of view are representative particle trajectories in yellow. Initially spherical particles were elongated into ellipsoids with a strain based on the projected 2D ellipse axes. **(B)** Fluid streaklines illustrated by fluorescent 1-µm tracer particles.

### Aqueous Methyl Cellulose

Aqueous methyl cellulose was used as the carrier fluid and buffer streams in particle image velocimetry and particle microfluidic creep experiments. Methyl cellulose has been used as a viscosifying agent in suspended cell microfluidic studies to deform cells using lower flow rates and low Reynolds number [5,27–29] and is known to not affect cellular biological functions [30]. Aqueous 0.85% w/v methyl cellulose was prepared by dissolving powdered methyl cellulose (Thermo Scientific Chemicals, CAS 9004-67-5, molecular weight 454.513 g/mol) in 80°C filtered DI water (30% of the final volume) with agitation and stirred for 30 minutes. Stirring is continued as the heating source was removed and the mixture was allowed to cool to 30°C. The remaining 70% of the water volume was chilled and added. The final mixture was stirred in an ice bath for 40 minutes. Methyl cellulose was filtered (0.2 µm PVDF filter, Cole Palmer) to remove debris and undissolved methyl cellulose. The viscosity was measured on a TA Instruments Discovery Hybrid series rheometer using a parallel plate geometry (40 mm diameter) with a shear rate sweep ramped logarithmically from 1 s^−1^ to 100 or 1000 s^−1^ with 10 points per decade. The viscosity used in the mechanical model was the average viscosity at the lowest reliable strain rates of 10 and 12.6 s^-1^ from 3 independent measurements.

*Fluid Theoretical Background.* The cross-slot microfluidic device aims to generate linear planar extensional flow with the velocity field given by

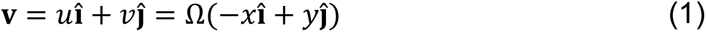

where the extensional strain rate Ω = −*მu*⁄*მx* = *მv*⁄*მy* is constant and homogeneous and the origin **X** = **0** is a point of zero velocity (stagnation point). In ideal planar extensional flow, the velocity does not depend on *z*. The velocity gradient ∇**v** is constant with diagonal non-zero terms equal to the extensional strain rate −*მu*⁄*მx* = *მv*⁄*მy* = Ω and zero off-diagonal terms *მu*⁄*მy* = *მv*⁄*მx* = 0. The velocity magnitude is proportional to the distance *r* from the stagnation point:

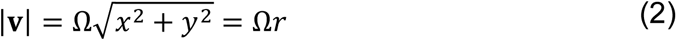

In the fluid field measurements, the presence of extensional flow was evaluated from the agreement of the measured velocity gradient with the ideal constant gradient (−*მu*⁄*მx* = *მv*⁄*მy* = Ω and *მu*⁄*მy* = *მv*⁄*მx* = 0), and the extensional strain rate was measured from Equation (1).

Equations that relate the strain rate to the microparticle trajectory points (*x*(*t*), *y*(*t*)) as it flows through the extensional flow region were found from integration of the velocity components *u* = *მx*(*t*)⁄*მt* = −Ω*x*(*t*) and *v* = *მy*(*t*)⁄*მt* = Ω*y*(*t*) along the streamline from a starting point (*x*(*t*_0_), *y*(*t*_0_)) = (*x*_0_, *y*_0_) at time *t*_0_ to a later point (*x*(*t*), *y*(*t*)) at *t*:

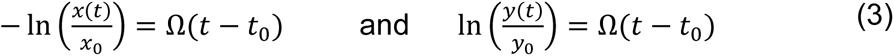

Combining to eliminate *t* and taking *e* to the power of both sides, the relationship between trajectory coordinates *x*(*t*) and *y*(*t*) is a right hyperbola (perpendicular asymptotes) with its center at the stagnation point located at (*x*_*SP*_, *y*_*SP*_):

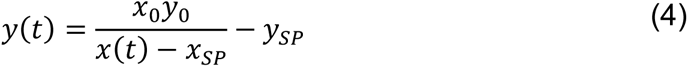

The location of the flow field’s stagnation point was estimated by fitting the particle trajectory (*x*(*t*), *y*(*t*)) to Equation (4) after picking an arbitrary point along the hyperbolic trajectory (*x*_0_, *y*_0_).

### Velocity Field Measurements

The unperturbed velocity field in the extensional flow device was measured using micro-particle image velocimetry (PIV) [32,33]. A suspension of 0.05% v/v 1-µm fluorescent polystyrene spheres (FluoSpheres, excitation/emission 540/560 nm, Molecular Probes) in aqueous 0.85% w/v methyl cellulose was infused into both device inlets using syringe pumps (Pump33 DDS and Pump11 Elite, Harvard Apparatus). Fluorescence microscopy movies of the flowing fluid were captured with a sensitive sCMOS camera at its fastest acquisition speed (Prime95B, Photometrics, full frame resolution 1200 × 1200 pixels, 12-bit depth, 1 ms exposure time, 81 frames per second, 12.4 ms between frames) at 10x magnification (NA 0.40, 0.9346 pixels per µm) and 40x magnification (NA 0.65, 3.759 pixels per µm) on an inverted microscope (Zeiss Axiovert 200M, Excelitas X-Cite 120LED Boost illumination system, Olympus TRITC 543 nm bandpass filter). The camera and motorized microscope were computer-controlled with MicroManager [34]. The velocity field was calculated from the movies with PIVlab v2.62 [35] using included image preprocessing and postprocessing vector validation features. Images were preprocessed with contrast limited adaptive histogram equalization (window size 20 px) [36] and a high pass filter (window size 15 px). Movie frames were sampled using three passes of interrogation areas with 50% overlaps (90 × 90, 44 × 44, and 22 × 22 pixels for 10x and 128 × 128, 64 × 64, 32 × 32 for 40x). Analyses with larger and smaller interrogation windows were performed to confirm convergence of the velocity answer. The pairwise correlation matrices within interrogation areas were evaluated in the frequency domain, and the peak locations of the cross-correlation functions are refined with subpixel precision using a 2D Gaussian function (3 px × 3 px neighborhood). Velocity vector outliers were identified with the normalized median check [37] (threshold 3) and global standard deviation check (threshold 8) and discarded. Missing vectors were replaced with values interpolated from nearest neighbors (3 px × 3 px neighborhood). Lower flow rates than particle creep experiments were used for PIV experiments due to the speed limitations of the sCMOS camera. Flow rates were chosen so that the maximum displacement of a particle was less than ¼ of the maximum interrogation area, a general PIV guideline [32].

Two kinds of PIV experiments were conducted to study the flow field characteristics. The first experiment focused on determining the flow regimes throughout the device and the dependence of fluid velocities and velocity gradient magnitudes with flow rate. The field of view at 10x magnification (1284.11 µm × 1284.11 µm, **Figure 1B**) captured all flow regimes: the fully developed Poiseuille flow in the inlet and outlet channels, the extensional flow in the central cross-slot region, and the transition region in between.

The fluorescent particle suspension was infused at several flow rates (50, 75, 100, 125 and 150 µL/hr), and 200-frame movies were captured at the device midchannel height (*z* direction parallel to gravity). Three technical replicates were captured per flow fate. The second PIV experiment explored the velocity field and gradient variations in the *z*-direction, thereby assessing how close the experiments matched the mechanical model’s assumption of planar extensional flow. These experiments were captured in 40-frame movies at various depths in 10 µm increments using a constant flow rate of 100 µL/hr with 40x magnification (40x has a thinner illumination volume than 10x). The field of view at 40x was the central extensional flow region (319.21 µm × 319.21 µm, **Figure S5**). For all movies, the reported velocity field was taken to be the average of the velocity vectors for each interrogation area across the frame pairs from the 200- or 40-frame movie.

Further post-processing of velocity vector fields was done to determine velocity gradients and measure the constant extensional strain rate in the central cross-slot region using a custom program in MATLAB (MathWorks, R2024a). The velocity gradient components were calculated using the central finite difference approximation for interior points and single-sided differences for edge points. The stagnation point location was calculated by fitting a paraboloid (Equation 2) to a 9 × 9 neighborhood centered at the location of the minimum measured velocity. The extensional strain rate in the central cross-slot region was measured from linear regression of the measured velocity magnitudes on their distance from the stagnation point for a circular region *r* ≤ 100 µm from the paraboloid-fitted stagnation point. This distance of 100 µm was chosen based on the consistency of standard deviation of velocity magnitude (**Figure S3, S4**), which indicated flow that closely conformed to the ideal uniform extensional flow. The unidirectional flows in the straight inlet and outlet channels of the cross-slot were compared to theoretical predictions of pressure-driven Poiseuille flow in rectangular channels [31]. *u*(*y*) for inlet channels and *v*(*x*) for outlet channels velocity profiles were averaged along the channel length up to 450 µm away from the stagnation point and fitted to parabolas (**Figure S7**). 450 μm from the stagnation point was sufficiently upstream of the transition zone to be pure Poiseuille flow.

### Agarose Microparticles

Agarose hydrogel microparticles were fabricated using an emulsion technique. 0.5% w/w aqueous agarose was prepared by adding agarose powder (Thermo Fisher Chemicals, CAS 39346-84-1) to filtered DI water and heating in an oven for 10 minutes. Mineral oil with 2% v/v Span 85 surfactant (Sigma Aldrich, CAS 26266-58-0) was heated to 70°C on a stirring hot plate. One millimeter of aqueous agarose was dispensed drop-wise with an 23-gauge stainless steel blunt needle into the 5 mL hot mineral oil bath under agitation (1250 rpm). Stirring continued for 4 more minutes, and then the emulsion was removed from heat and allowed to cool to room temperature. After overnight settling, the agarose hydrogels were washed using DI water over 4 rounds of centrifugation and oil supernatant removal. The concentrated agarose particle suspension (the washed particles recovered from the original 1 mL of 0.5% w/w aqueous agarose) were stored at 4°C. The agarose hydrogels were spherical. Most particles analyzed in the microfluidic creep experiments had diameter 25 – 50 μm, and a few larger particles 80 – 85 µm were analyzed (**Figure 6A, 6A**).

### Microfluidic Creep Experiments

To measure linear viscoelastic mechanical properties, agarose microparticles were observed flowing in the straight inlet channel and stretching in the extensional flow region until the particle left the field of view by way of one of the 2 outlets (**Figure 1A**). The agarose microparticles suspension was prepared by distributing 250 µL of concentrated agarose particles in 5 mL aqueous 0.85% (w/v) aqueous methyl cellulose. Particles were infused into only one inlet with blank 0.85% w/v aqueous methyl cellulose infused in the opposing inlet. The total flow rates through each inlet were 6.5, 7.5, 8.5, and 9.5 mL/hr with a sheath-to-center stream flow rate ratios of 2.2 - 2.25. These ratios resulted in a center stream width (65 µm) that was slightly larger than most particles measured in this study (25 – 50 µm diameters). The fluid Reynolds numbers were small and therefore inertia could be reasonably neglected: 0.061, 0.070, 0.079, and 0.088 using viscosity *µ* ∼ 110 mPa·s from our shear rheology measurements, fluid density *ρ* ∼ 1065 kg/m^3^ [28], average fluid velocity in straight channel *U = Q/(hw*), and channel dimensions *w* = 290 µm and *h* = 265 µm. Three-second particle deformation movies were captured under phase contrast microscopy at 10x magnification (Zeiss Axiovert 200M inverted microscope, 0. 9983 pixels per µm) using a high-speed camera (JetCam 19M and Komodo Frame Grabber, Kaya Instruments). The microfluidic system was equilibrated for at least 2 minutes after increasing the syringe pumps to a higher flow rate. Particle movies were captured at the device mid-height so that in-focus particles were at approximately the center of the device and as far from the channel walls as possible. Frames per second varied with flow rate to reduce blurring at faster particle speeds: 3800 fps at 6.5 mL/hr, 4200 fps at 7.5 mL/hr, 4600 at 8.5 mL/hr, 5000 at 9.5 mL/hr. Exposure time was a constant 88 µs. Camera region of interest was 1200 px by 352 px and biased towards the particle suspension inlet channel to capture the incoming particle shape. A single batch of agarose microparticles was measured on two different days in the same microfluidic device and using the same aqueous methyl cellulose batch for the suspending fluid.

### Particle Experiment Image Analysis for Kinematics and Deformation

Microparticle trajectories and deformation were analyzed using a custom MATLAB analysis code with some open-source sub-functions. Single particles that were sufficiently in-focus with no nearby neighboring particles were analyzed for their trajectory through the device and then shape at each time point along the trajectory as follows. First the background was subtracted from all movie frames. The background was the time-average image of all movie frames. The background-subtracted movie frames were smoothed using a Gaussian filter (9 px × 9 px filter size, Gaussian distribution standard deviation 2 px) and binarized using a global threshold value that was manually selected prior to analysis to detect most agarose microparticles. The approximate particle shape and location was determined from objects in the binarized movie frame that had areas (500 ≤ *A ≤* 8000 px^2^) and circularities (0.9 ≤ *C =* 4π*A*/*L*^2^ ≤ 1.1 where *L* is the object perimeter) in the ranges expected for the 25-75-µm-diameter particles while all other objects were discarded. The object center of mass locations were linked into trajectories using the MATLAB adaptation [38] of a particle tracking algorithm [39,40]. Next, the edges (contours) of particles at each time point along their trajectories were found. The background-subtracted movie frames were smoothed using a Gaussian filter (2 px × 2 px filter size, Gaussian distribution standard deviation 0.5 px). The particle contour location was defined as the location of maximum grayscale intensity gradient along rays (360 total) drawn from the center of the particle. Sub-pixel edge detection resolution was achieved by finding the maximum of a parabola fit to the local intensity gradient in a 5-pixel neighborhood. Because the initially spherical particles were expected to stretch into ellipsoids, an ellipse was fitted to each particle contour using the least-squares criterion [41]. The particle center of mass locations were updated to the center of mass of the contour points for each frame. The particle strain was defined from the ellipse axes as the familiar Taylor deformation parameter: *ɛ* = (*b* − *a*)/(*b* + *a*) where *b* is the ellipse semi-axis in the vertical direction (parallel to *y* or outlet axis), and *a* is the ellipse axis in the horizontal direction (parallel to *x* or inlet axis) (**Figure 1A**). A positive strain meant the particle elongated in the *y* direction, a negative strain meant the particle stretched in the *x* direction, and zero strain meant the detected particle was an unstitched sphere. The equivalent particle radius in each frame was calculated from the ellipse axes, assuming that the spherical particle was slightly deformed into an ellipsoid with the out-of-plane semi-axis equal to the smaller of the in-plane *a* and *b* semi-axes: *r*_*p*_ = (*ab* · min(*a*, *b*))^1/3^. The no-flow equilibrium particle diameter *d*_0_ = 2*r*_0_ was estimated from *r*_*p*_ in the Poiseuille-to-extensional flow transition zone around 250 µm from the stagnation point where the particle experienced the lowest hydrodynamic forces. Particle trajectory arc length was the cumulative Euclidian distance of the particle center of mass trajectory points. Particle speed in each frame was estimated using the central difference approximation for interior points and forward and backward difference approximations for the first and last time points. Particle inlet speed was the average of particle velocities for trajectory points ≥ 450 µm from the stagnation point. Violin plots [42] were used to display particle parameter distributions.

In steady flow, the particle trajectory aligned with fluid streamlines, so the particle trajectory was used to estimate extensional flow field features. To determine the stagnation point location, the particle’s center of mass trajectory was fit to a right hyperbola, the expected path of an entrained particle in uniform extensional flow (**Equation 4**). The hyperbola was fit to trajectory points within 100 µm of the user-defined approximate stagnation point, where PIV measurements showed the extensional strain rate, i.e., the slope of |**v**| vs. *r*, was constant within measurement precision (**Figure 2, S4, S5**). Prior to fitting the right hyperbola to the particle trajectory in the constant extensional flow region, the slight rotation angle between the particle trajectory and the camera orientation was determined from the linear regression fit to the trajectory points in the inlet channel region where the particle traveled in a straight line. The angle was ≤ 1° for all movies.

**Figure 2.**
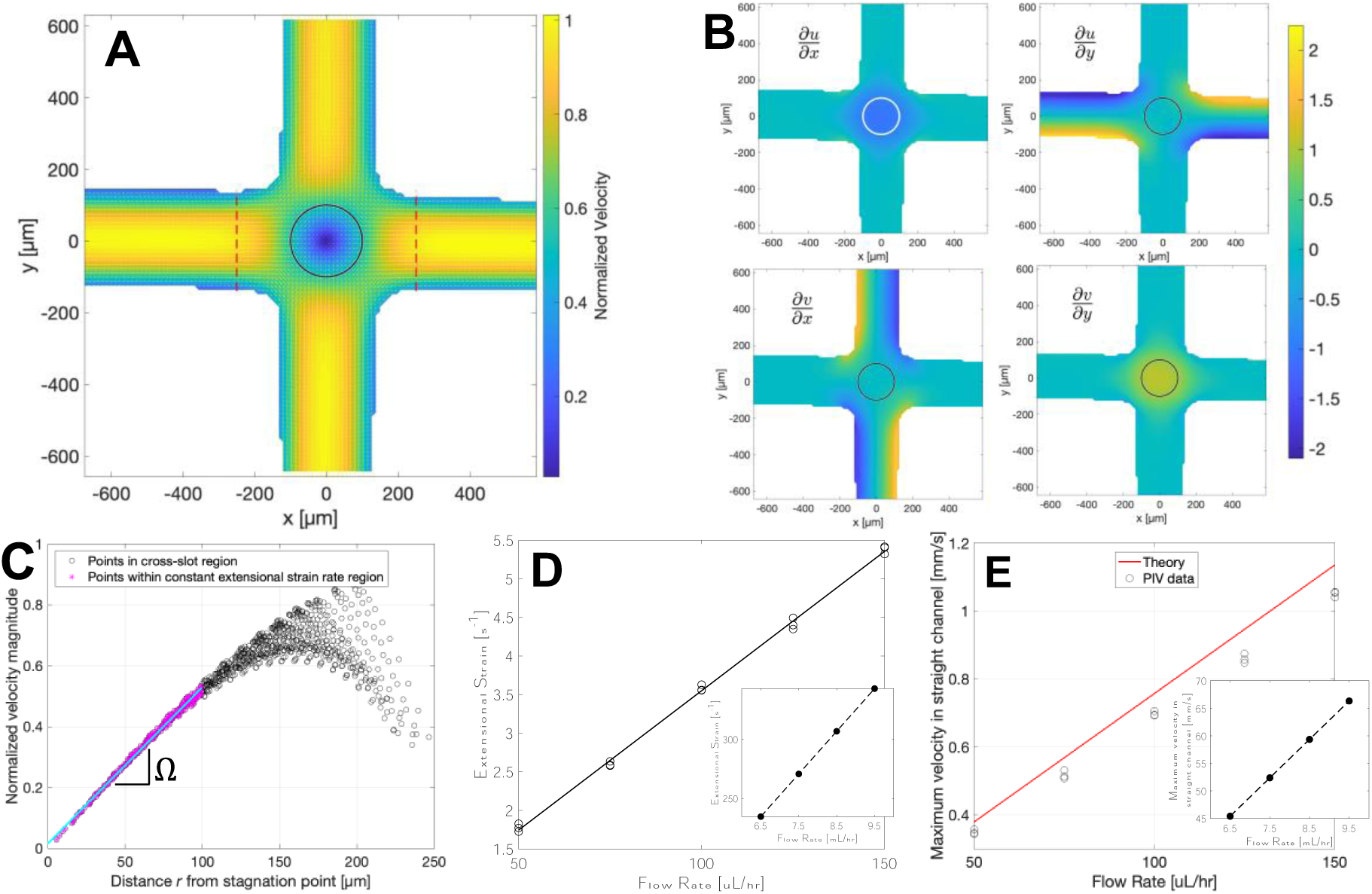
Particle image velocimetry measurements (PIV) of the cross-slot flow field. **(A)** Fluid velocities measured with PIV. The 100-µm-radius black circles indicate the region of constant extensional strain rate. The red dashed lines mark 250 µm upstream of the stagnation point where the extensional flow velocity gradient started, and particles began to feel extensional flow. The velocity was normalized by the maximum velocity from parabolic fits to the unidirectional velocities averaged over the 4 straight channels. **(B)** Velocity gradient components normalized by the constant extensional strain rate Ω in the central 100-um-radius circular region (circled area). The diagonal components were uniform in the central cross-slot region. The off-diagonal components showed the parabolic Poiseuille flow profiles in the straight channels and zero in the device. **(C)** The constant extensional strain rate Ω was the slope of the least-squares linear regression line for the velocity magnitude |**v**| with distance from the stagnation point *r*. Data in (A – C) are from a representative PIV experiment with 100 µL/hr flow rate infused to both inlet channels. **(D)** The constant extensional strain rate Ω from PIV experiments at 5 flow rates. Inset: Extrapolation of Ω to flow rates used in the agarose microparticle experiments. **(E)** The maximum velocity magnitude in the Poiseuille flow of the straight channels for PIV experiments at 5 flow rates compared to the analytical solution for pressure-driven flow through a rectangular channel [31]. The maximum velocity for each PIV data point is the average maximum velocity of the four straight channels, where in each channel, a parabola was fit the lengthwise-averaged velocity profiles for *r* ≥ 450 µm. Inset: Maximum velocity trend from PIV data extrapolated to flow rates used in particle experiments.

### No-flow Particle Shape

Hydrogel particle shape was analyzed in a fluid-filled chamber (CoverWell, 9 mm wide, 0.5 mm deep, Invitrogen) under no-flow conditions to assess the typical particle shape in the absence of deforming forces. Furthermore, these data provided information about the minimum measurable particle strain of our experimental and image processing techniques. The same imaging conditions (10x magnification, phase contrast microscopy), camera (JetCam 19 high-speed camera), and particle edge detection image processing algorithm from the microparticle microfluidic creep experiments were used. Particles were imaged at their equator in 250-frame movies (4 seconds elapsed time, 50 fps, 88.5 µs exposure time).

### Particle Mechanical Theory

In a traditional creep test, a stress is suddenly applied to an object, and its time-dependent strain response is recorded. In the microfluidic cross-slot, this loading condition is approximated as particles enter and dwell in the cross-slot device extensional flow region. Using the elastic-viscoelastic correspondence principle [43], the Laplace-transformed viscoelastic solution of a linear viscoelastic sphere deforming in planar extensional flow was obtained directly from the corresponding elastic solution. From Murata’s solutions for an incompressible elastic sphere deforming in linear flows of a Newtonian fluid at low Reynolds number [44], we obtained at the strain of an elastic sphere at its equator when deformed at the stagnation point of planar extensional flow:

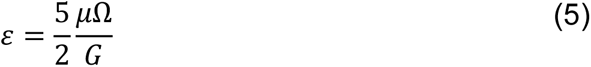

Ω is the uniform velocity field extensional strain rate, *μ* is the fluid viscosity, and *G* is the sphere’s shear elastic modulus. To arrive at the solution of the viscoelastic problem, the quantities in Equation (5) were reinterpreted as their Laplace transforms in the *s-*domain and the inverse Laplace transform was applied to arrive at the time-dependent viscoelastic solution. Assuming the agarose microparticles can be approximated by the Kelvin-Voigt constitutive model for viscoelastic solids (spring and damper in parallel), the time-dependent strain for a viscoelastic sphere in suddenly applied planar extensional flow is:

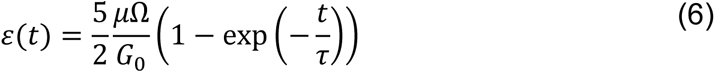

*G*_0_ is the elastic part of the mechanical response, *τ* is the retardation time, and *η* = *G*_0_*τ* is the viscous part of the response. More derivation details are available in the Supporting Information.

## RESULTS

PIV measurements indicated that the cross-slot device generated uniform planar extensional flow in a 100-µm radius region centered at the stagnation point. The PIV measurements confirmed the expected Poiseuille flow in the inlet and outlet channels and the low-speed region where the two straight channels intersect (**Figure 2A**). The extensional diagonal components of the velocity gradient *მu*/*მx* and *მv*/*მy* were very uniform in the cross-slot intersection and practically zero in the straight channels (**Figure 2B**). Furthermore, *მu*/*მx* and *მv*/*მy* were equal in magnitude and opposite in sign, as expected for planar extensional flow. The off-diagonal velocity gradient components *მu*/*მy* and *მv*/*მx* showed linear cross-channel behavior in the inlet and outlet channels, respectively, indicating parabolic Poiseuille flow profiles in these regions (**Figure 2B**). Shear velocity gradient components were also high at the device corners and very low to zero elsewhere.

The region of constant extensional flow was found by inspecting how far from the stagnation point the velocity magnitude vs. distance scatter plot was linear with roughly constant variation (**Figure 2C**), an indication that the fluid flow was conforming to extensional flow (Equation (1)). Across the five flow rates and using 10x magnification, this distance was found to be about 100 µm for this device (**Table S2**). The constant extensional strain rate Ω was estimated from the slope of the linear regression line within this 100 µm radius region for all flow rates (**Figure 2C**) and increased proportionally with flow rate (**Figure 2D**). The flow in the straight inlet channels had a parabolic profile with flow in only one direction (**Figure S7**), as expected for Poiseuille flow. The average maximum flow rate of all four straight channels increased linearly with flow rate and was less than 10% below the value predicted from theory for a 290-µm-wide, 265-µm-deep channel at the same flow rates [31] (**Figure 2E**). Given the presence of volume illumination during PIV experiments and channel dimension uncertainty due to wall imperfections, this level of agreement between theory and experiment was reasonable. The linear fits to the measured constant extensional strain rate and maximum inlet/outlet channel speeds were extrapolated to the higher flow for comparison with hydrogel particle experiment results (insets in **Figure 2D, 2E**). At 40x magnification, the constant extensional strain rate was very constant ± 20 µm above and below the channel midheight (**Figure S5)**. Taken together, the PIV data indicated that the undisturbed flow closely matched the uniform planar extensional flow assumed in the linear mechanical model in a large 100-µm radius region around the stagnation point.

The 0.85% w/v aqueous methyl cellulose viscosity used in in the mechanical model Equation (6) is taken to be the fluid’s zero-shear viscosity. We estimated the zero-shear viscosity to be *μ* ∼ 110 mPa·s from the average viscosity at low shear rates (**Figure S1**, **Table S1**). We assumed the effective fluid viscosity felt by the deforming particles would be close the fluid’s zero-shear viscosity because of the low shear rates (off-diagonal velocity gradient terms *მu*⁄*მy* , *მv*⁄*მx*) in the extensional flow region measured in PIV experiments (**Figure 2B**, **Figure S6**).

Flow-focused particles were stretched in the extensional flow and achieved a quasi-steady strain state under suitable conditions. The sheath-to-center-stream flow rate ratio of 2.25 created a center stream width of ∼65 µm immediately upstream of the extensional flow region and focused most particles, typically less than 50 µm in diameter, near the center streamline (**Figure 1A**). **Figure 3** illustrates a representative microparticle experiment. Particles approached the stagnation point and followed a hyperbolic trajectory out the top or bottom inlet after dwelling in the constant extensional flow region for up to tens of milliseconds (**Figure 3A**). In the straight inlet channel, particles were measured to have small negative strain, meaning they were measured to elongated in the incoming flow direction (timepoint 1); the magnitude of these incoming strains, typically around -0.01, which is above the average strain of 0.006 ± 0.003 (mean ± standard deviation, *N* = 36) for particles in a still fluid chamber (**Figure S2**).

**Figure 3.**
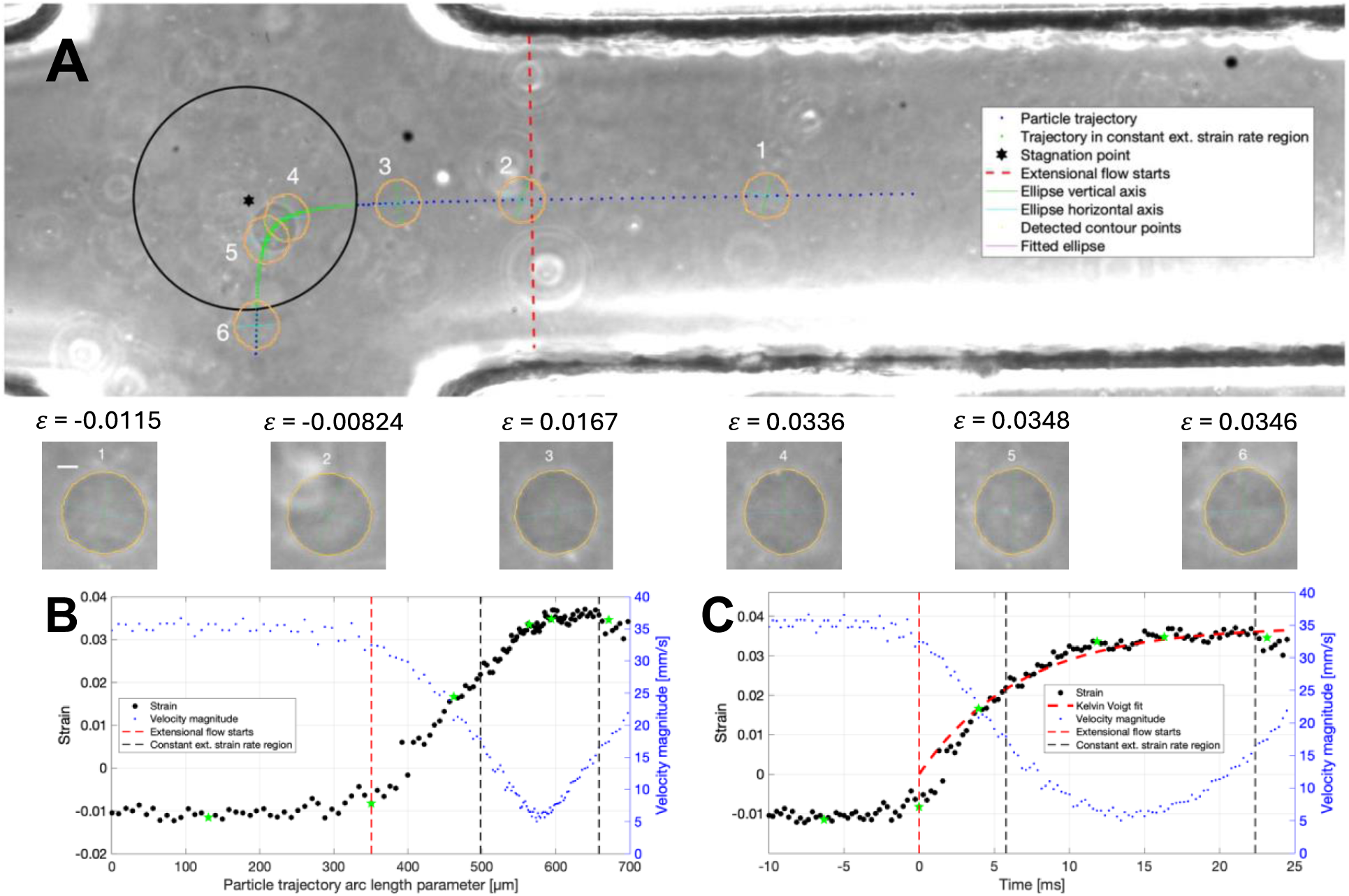
A representative microfluidic creep test result for 0.5% w/v agarose microparticles. **(A)** A 40.6-µm-diameter agarose particle’s trajectory (blue dots) through the microfluidic device, with extensional flow becoming significant at the red dashed line. The green trajectory points within the 100-um-raidus constant extensional strain rate region (black circle) were fit to a right hyperbola to estimate the stagnation point location (black star). Zoomed images of the particle’s detected contour and the best-fit ellipse along with the ellipse’s strain are shown for 6 time points. Scale bar is 10 µm. The particle strain (left axis) and speed (right axis) are plotted against **(B)** the particle trajectory length and **(C)** time. The particle started elongating away from its incoming strain level at 250 µm upstream of the stagnation point (red dashed vertical line). As the particle dwelled at low speed in the constant extensional strain rate region (between black dashed vertical lines), the particle reached a quasi-steady strain state. In (C), fitting Equation (6) (curved red dashed line) yielded Kelvin-Voigt parameters of *G*_0_ = 1723 Pa, *τ* = 6.7 ms, and *η* = 11.6 Pa·s. The green stars mark the locations of time points 1-6 from (A). The flow rate for these representative data was 6.5 mL/hr total into each inlet. From extrapolated PIV results, the fluid extensional strain rate was 234.59 s^-1^.

Approximately 250 µm upstream of the stagnation point, the particles began to slow down and elongate in the outlet flow direction (timepoints 2-3). Well-centered particles could remain stretched at a steady strain until leaving the camera field of view out an outlet channels (timepoints 4-6). The 40.6-µm particle in **Figure 3** reached a minimum distance of 32.5 µm from the stagnation point where it slowed down to a minimum speed of 5.0 mm/s.

Particle position, within the device or along its trajectory arc length, was an important independent variable for analyzing particle deformation. Like the position-focused analysis approach of unidirectional extensional flow techniques with droplets [45,46] and cells [29], the particle’s location within the cross-slot extensional flow device determined the hydrodynamic fluid forces deforming it and its deformation state. Plotting particle strain against trajectory arc length, points crowded in the latter part of the trajectory in the constant extensional flow region where the particle did not translate far between movie frames (**Figure 3B**). This slow particle motion as it dwelled near the stagnation point corresponded to over 10 ms of elapsed time in the constant extensional flow region (**Figure 3C**). For the 0.5% w/v agarose microparticles, this amount of time was sufficient for many particles to achieve a quasi-steady stretched state. The **Figure 3** particle spent 16.6 ms in the constant extensional strain rate region and stayed at a steady strain of 0.035 for approximately 5 ms before leaving the region.

The particle trajectory data was correlated with the PIV fluid field measurements to gain insight into the hydrodynamic forces causing the particle deformations observed in Figure 3. The particle’s speed followed the fluid velocity trends of the fluid **(Figure 4A),** though the particle’s average inlet speed (35.35 mm/s) was 22% slower than the maximum fluid speed predicted from extrapolated PIV data (45.4 mm/s). The plots of velocity gradient components interpolated onto the particle trajectory points showed the underlying reason why the particle began elongating around 250 µm upstream of the stagnation point (**Figure 3B, 3C**): at this location, the diagonal components of the velocity gradient have increased significantly from about zero to 20% of Ω, the constant extensional strain rate in the 100-µm-radius region around the stagnation point (**Figure 4B, 4C**). At this point, the particle was responding to the extensional flow by elongating in the outlet flow direction. The representative 40.6-µm-diameter particle from Figure 3 and Figure 4 was in the constant extensional strain rate region only 46% of the length of the 347-µm-long trajectory section on which it experienced some extensional flow (| *მu*⁄*მx*| , |*მv*⁄*მy*| > 0). Yet the particle spent the majority of the total 24.5 ms along this trajectory section, 16.6 ms or 68% of the time, in the uniform extensional strain rate region. Thus, application of a model to extract mechanical properties that assumes the particle experiences constant extensional flow was a reasonable approximation.

**Figure 4.**
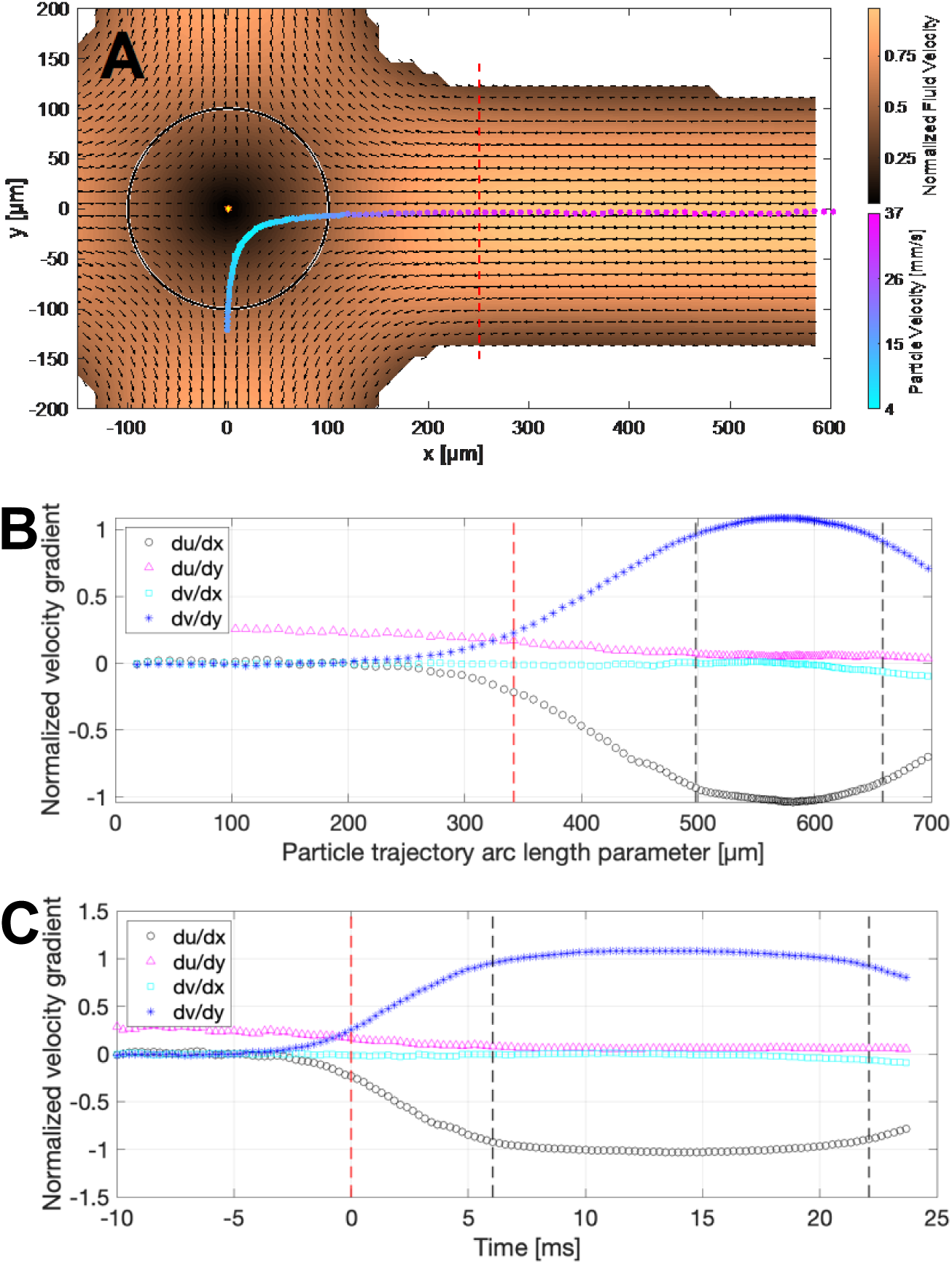
Correlating microparticle location with the fluid velocity and velocity gradient field along its trajectory**. (A)** The trajectory points and speed of the representative 40.6-µm-diameter particle were overlayed on the measured fluid field from a particle image velocimetry (PIV) experiment without the particle present. The fluid velocity was normalized by average parabola-fitted maximum fluid speed in the four straight channels. The PIV velocity gradient was interpolated at the particle’s trajectory points and plotted against **(B)** the particle trajectory length and **(C)** time. The particle experienced minimal deforming velocity gradient forces up until 250 µm upstream of the stagnation point (red dashed vertical line). At this point, the extensional diagonal components of the velocity gradient increased until plateauing in constant strain rate 100-µm-radius region (between the black dashed vertical lines). The off-diagonal gradient components remain low along the trajectory. The velocity gradient components were normalized to the constant extensional strain rate Ω. The PIV data is from the representative PIV experiment with 100 µL/hr flow rate shown in Figure 2.

The uniaxial microscale creep test took place while the particles felt the extensional flow. We determined 250 µm upstream of the stagnation point, a distance slightly less than the channel width of 290 µm, to be a reasonable location where particles begin to experience extensional flow. From the undisturbed fluid PIV measurements, the velocity gradient near the central streamlines where particles would be located changed from approximately zero to Ω (**Figure S6**). Though the diagonal components *მu*/*მx* and *მv*/*მy* rose above the noise level close to 350 µm upstream of the stagnation point, particles didn’t elongate immediately, and the extensional flow needed to get strong enough to start deforming the particle. We chose 250 µm after looking at 140 particles and observing which choice led to the best collapse of the strain data when plotting against particle trajectory arc length *s* and setting *s* = 0 to where extensional flow started for each particle (**Figures S12 – S15**). Because particles were still experiencing extensional strain rates close to the constant strain rate Ω when leaving the camera field of view (**Figure 4B**, **Figure 4C**), the mechanical model Equation (6) was applied from 250 µm upstream of the stagnation point to the time point when the particle left the field of view. The mechanical model yielded reasonable fits to the strain data (**Figure 3C**); for this representative particle, the fitted Kelvin-Voigt parameters were *G*_0_ = 1723 Pa, *τ* = 6.7 ms, and *η* = 11.6 Pa·s.

Not all particles were sufficiently centered to dwell in the extensional flow region long enough to achieve the quasi-steady stretched state. Well-centered particles looked visually very centered in the straight inlet channel and closely approached the stagnation point (**Figure 5A**). This close proximity to the stagnation point and particle velocity slowdowns allowed the particles to dwell long enough >15 ms to reach steady strain, and many particles experienced a decrease in strain after leaving the uniform extensional strain rate region (**Figure 5B**). Other particles were visibly off-center in the inlet channel and did not pass close to the stagnation point (**Figure 5C**). The strain of these particles was still increasing when the particles left this region after only ∼10 ms (**Figure 5D**). In this small example data set, the well-centered particles tended to be bigger (mean ± standard deviation 40.2 ± 9.4 µm, *N* = 9) than the poorly centered particles (31.1 ± 4.6 µm, *N* = 11) at the flow rate ratios used. The offsets to the centerline upon reaching the constant extensional strain rate region (*r* ≤ 100 µm) were over five times larger for the poorly aligned particles (22.8 ± 9.4 µm) compared to well-centered particles (4.63 ± 2.9 µm). The frequency of incoming particles that were well-centered and poorly centered were approximately equal at the flow rates and particle concentration tested: 1 – 2 well-centered particles per second flowed through the cross-slot device along with 1 – 2 poorly centered particles per second. The fitted Kelvin-Voigt elastic parameters for both cases were similar: 1650 ± 171 Pa for the plateaued strain particles compared to 1641 ± 189 Pa for the particles lacking a strain plateau (**Figure S9**). However, the edge detection algorithm struggled with the low-contrast agarose particles, and more accurate data would be needed to make a strong claim about the level of inaccuracy in mechanical property measurements for non-plateauing particle. The viscous part of the two particle alignment cases showed greater discrepancy: the poorly aligned particles that did not reach a strain plateau were measured to be more viscous (12.3 ± 3.4 Pa·s) compared to the more centered particles that had a quasi-steady strain state (9.4 ± 1.5 Pa·s) (**Figure S9**).

**Figure 5.**
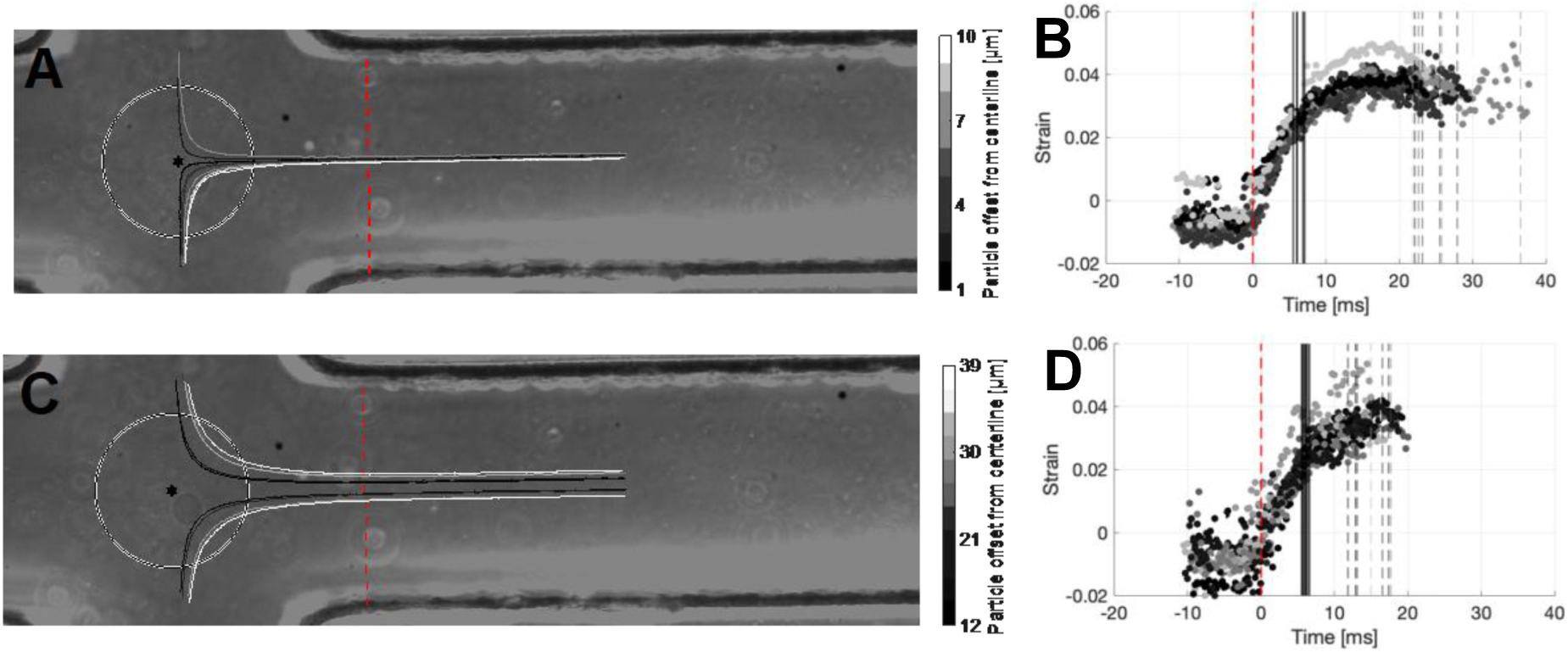
Comparing the trajectories and resulting particle strains for representative well-centered 0.5% w/v agarose microparticles and poorly aligned particles relative to the stagnation point for a flow rate of 6.5 mL/hr. (**A)** Well-centered particles followed a high curvature trajectory that closely approached the stagnation point. *N =* 9 particles. **(B)** The centered particles dwell in the stagnation point region for 15+ ms and achieved a strain plateau that lasted several milliseconds. **(C)** Poorly centered particles did not get as close to the stagnation point and followed a lower curvature hyperbolic trajectory. *N =* 11 particles. **(D)** The poorly aligned particles dwelled for a shorter time in the constant strain rate region (∼10 ms) and did not achieve a strain plateau. The strain was still increasing when these particles left the constant strain rate region. For all subfigures, the grayscale trajectory and strain marker colors correspond to the particle’s offset from the center streamline upon entering the constant extensional strain rate region (black circle). In (B) and (D), the particle strain data were well aligned by choosing *t* = 0 to correspond to the location 250 µm upstream of the stagnation point where the extensional flow starts (red dashed vertical line). The solid black vertical line marks when the particles entered the constant extensional strain region (*r* ≤ 100 µm), and the grayscale dashed vertical line marks where individual particles left this region. The flow rate for these representative data was 6.5 mL/hr total into each inlet. From extrapolated PIV results, the fluid extensional strain rate was 234.59 s^-1^.

**Figure 6.**
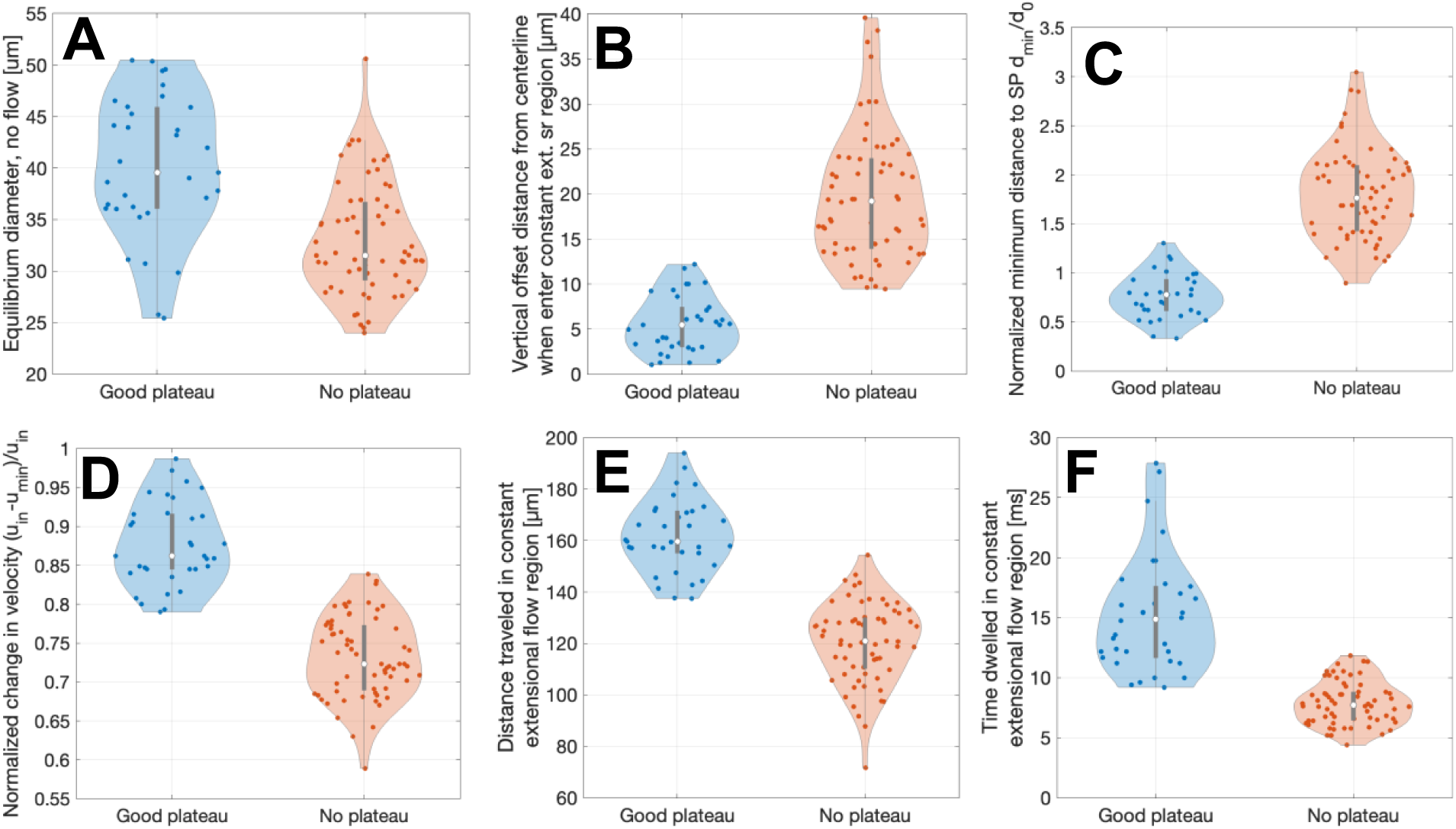
Distributions on 0.5% w/v agarose microparticles kinematics for particles that achieved a strain plateau in the extensional flow cross-slot region (‘Good plateau’) and particles that did not achieve a plateau whose strain was still increasing when leaving the camera field of view (‘No plateau’) from all four flow rates used. **(A)** Estimates of particle diameter in no-flow conditions estimated from its projected area at the location 250 µm upstream of the stagnation point. **(B)** The particle offset from the center streamline, here measured when the particle entered the constant extensional strain rate region (≤100 µm from stagnation point). **(C)** The minimum distance from the particle to the stagnation point along its trajectory normalized by the particle diameter. **(D)** The change in particle speed from the straight inlet channel to its minimum speed near the stagnation point normalized by the inlet speed. **(E)** Distance traveled by particles in the constant extensional strain rate region. **(F)** Time particles spent in the constant extensional strain rate region. Violin plots show the kernel density estimate of the data, individual data points overlay, and a box-and-whisker plot showing the median (white circle), quartiles (box), and non-outlier range (whiskers) with outliers defined as 1.5 × interquartile range (box height). Number of particles measured plateau condition combining experiments from all four flow rates (6.5, 7.5, 8.5, 9.5 mL/hr): Good plateau *N* = 34, No plateau *N* = 64.

Differences in kinematic parameters between 0.5% w/v agarose microparticles that achieved a strain plateau in the constant extensional strain rate region and those that did not were investigated. The motivation was to explore easily measurable criteria related to particle position and speed that can be used to select for plateaued particles that were situatable for mechanical property measurement. The plateaued and non-plateaued particles came from a similar size range 25 – 50 µm diameter (**Figure 6A**). The kinematic parameters in **Figure 6B**-**Figure 6F** were well separated between strain plateaued and non-plateaued particles with little overlap. The particle offset from the center streamline, here measured when the particle entered the constant extensional strain rate region (≤100 µm from stagnation point), was the initial condition that determined downstream metrics. The plateaued particles were offset no more than 12 µm while the particles without a strain plateau had offsets higher than 9.5 µm (**Figure 6B**). Near the stagnation point, plateaued particles tended to be within a particle diameter of the stagnation point, while non plateaued particles were generally further than a particle diameter away (**Figure 6C**). The change in normalized particle velocity between the straight inlet channel and the minimum speed at the stagnation point showed a transition point around 0.8 between the two groups (**Figure 6D**). Within the extensional flow region, strain plateaued particles traveled ∼140+ µm over ∼10+ ms, and non-plateaued particles typically spent less time in that region (**Figure 6E**, **Figure 6F**).

We studied strain-plateaued particles deforming four flow rates to investigate the impact of flow rate on particle trajectories and the microfluidic creep test results. The no-flow particle diameters were mostly in the range 25 – 55 µm and a few in the size range 80 – 85 µm (**Figure 7A**). The maximum particle speed increased linearly with flow rate, as expected, for all particles including the 3 particles larger than 55 µm. The particle speeds were about 20% lower than the predicted fluid speed from PIV (**Figure 7B**). On the inlet channel Poiseuille flow, most particles were measured to have an average strain that was negative, meaning the particles were elongated in the inlet flow direction, and the magnitude of this strain increased slightly with flow rate (**Figure 7C**). However, for many particles the edge detection algorithm picked up many inaccurate edge sections over portions of the particle contour as the low-contrast, fast-moving agarose which were traversing a non-uniform background with small fluid or device inhomogeneities creating large refractive index variation that was amplified by phase contrast microscopy. There can be large variation in the detected incoming particle shape (**Figure S12 – S15**), so the data trends in **Figure 7C** need more investigation before drawing conclusions about particle incoming shape. The edge detection algorithm performed well as the particle slowed down and entered extensional flow, and we have higher confidence in the particle contour-derived data from the central cross-slot region of the device. The particle offset distributions had large overlap across flow rates with a slight increase with flow rate (**Figure 7D**). The minimum particle distances to the stagnation point did not significantly change with flow rate either (**Figure 7E**). The average and range of minimum particle speeds increased but there was a lot of overlap for all flow rates (**Figure 7F**).

**Figure 7.**
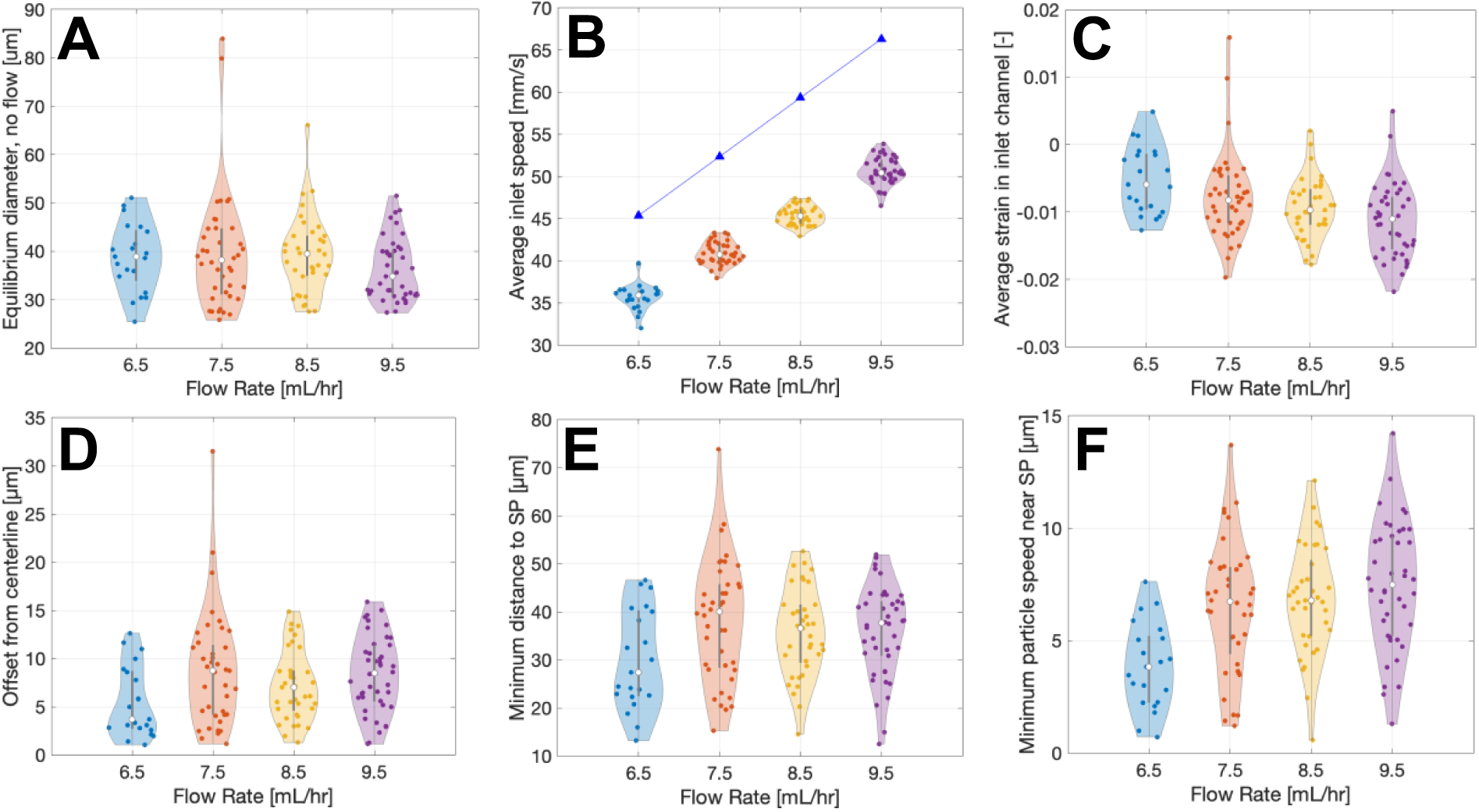
Distributions on 0.5% w/v agarose microparticles kinematics for the incoming straight channel and extensional flow region in the cross-slot microfluidic device for four flow rates. **(A)** The particle equilibrium diameter in no-flow conditions estimated from its projected area at 250 µm upstream of the stagnation point. **(B)** Particle speed in the straight inlet channel averaged over trajectory points ≥ 450 µm upstream of the stagnation point. The blue diamonds are the predicted maximum fluid velocity extrapolated from particle image velocimetry experiments. **(C)** Average particle strain in the inlet channel. **(D)** Particle offset from the center streamline when entering the 100-µm-radius constant extensional strain rate region. **(E)** Minimum distance of the particles to the stagnation point. **(F)** Minimum particle speed near the stagnation point. Violin plots show the kernel density estimate of the data, individual data points overlay, and a box-and-whisker plot showing the median (white circle), quartiles (box), and non-outlier range (whiskers) with outliers defined as 1.5 × interquartile range (box height). Number of particles measured per flow rate: 6.5 mL/hr: *N* = 21, 7.5 mL/hr: *N* = 40, 8.5 mL/hr: *N* = 37, 9.5 mL/hr: *N* = 42. All particles achieved a strain plateau.

The mechanical model from Equation (6) was applied to obtain the Kelvin-Voigt material parameters for the strain plateaued particles at the four flow rates used in microfluidic experiments (**Figure 8**). We observed an increasing trend in shear modulus *G*_0_ and a slight decreasing trend in retardation time *τ* with increasing flow rate. This made for a relatively consistent viscosity *η* = *G*_0_*τ* across flow rates. However, many particles had appreciable variation in the strain plateau due to debris or inhomogeneities being detected over the low-contrast agarose particle boundary, so the distributions of the material properties were large. Furthermore, these particles were made using a chaotic emulsion technique, so some of the measured material property variation could be due to physical inter-particle variation (different agarose and/or cross-linking densities).

**Figure 8.**
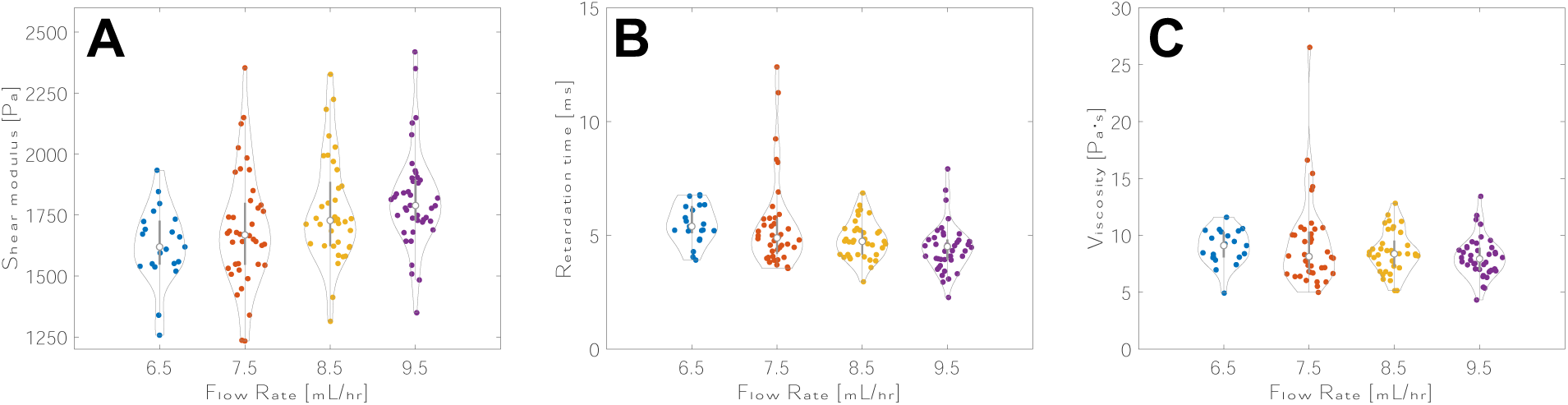
Fitted Kelvin-Voigt parameters for 0.5% w/v agarose microparticles in the microfluidic creep test using the extensional flow cross-slot device. The distributions of **(A)** the elastic stiffness *G*_0_, **(B)** the retardation time *τ*, and **(C)** viscosity *η* = *G*_0_*τ* are shown for four flow rates corresponding to the following extensional strain rates extrapolated from particle image velocimetry results: *Ω* = 234.6, 270.7, 306.8, and 342.9 s^-1^ for flow rates 6.5, 7.5, 8.5, and 9.5 mL/hr. Violin plots show the kernel density estimate of the data, individual data points overlay, and a box-and-whisker plot showing the median (white circle), quartiles (box), and non-outlier range (whiskers) with outliers defined as 1.5 × interquartile range (box height). Number of particles measured per flow rate: 6.5 mL/hr: *N* = 21, 7.5 mL/hr: *N* = 40, 8.5 mL/hr: *N* = 37, 9.5 mL/hr: *N* = 42. All particles achieved a strain plateau.

## DISCUSSION

We have demonstrated the ability of the cross-slot microfluidic device to approximate a traditional creep test at relatively high-throughput for microscale spherical objects. Microparticles sufficiently close to the central streamline rapidly experienced extensional flow within a few milliseconds and then reached a quasi-steady strain plateau while dwelling in the constant extensional strain rate region near the stagnation point for over 10 ms. This strain plateau will be key for accurately and precisely measuring linear viscoelastic properties of small microscale biological objects. This mechanical test was performed in the linear regime, with low Reynolds numbers Re < 0.1 and low particle strains *ɛ* < 0.1, where linear viscoelasticity theory and Stokes flow applied. With channel dimensions relatively large compared to particle diameters, an analytical mechanical model was proposed that was derived from the linear elasticity solution using the elastic-viscoelastic correspondence principle.

The numerical study of Lu et al. [22] predicted the quasi-steady strain state of particles passing through a cross-slot device, but we are not aware of an experimental study reporting how to achieve this plateaued strain state. The size discrepancy between the channel width and particle size was an important design parameter for reliably achieving a quasi-steady strain state in experiments by setting the particle dwell time in the extensional flow region, along with the flow rate. We observed roughly 25% of in-focus particles obtaining a quasi-steady deformation state as they dwelled near the stagnation point when the channel width (290 µm) was 6 – 12 times greater than particle size (mostly 25 – 50 µm diameters). Most of the particles’ time in extensional strain was spent in the constant extensional strain rate region (*r* ≤ 100 µm), which was a circular area with radius roughly 1/3 of channel width and 4 – 8 times larger radius than the suspended particles. Previous cross-slot studies that did not report a quasi-steady strain state used channel dimensions that were closer to cell size. Low Reynolds number experimental studies performed on cells 10 – 20 µm in diameter used channels that were 1.5 – 7 times wider than the cell diameters. Data from Armistead et al. [5] showed that cell strain was still increasing when cells left the central cross-slot region after 5 ms in a device that was 35 µm wide and 25 µm deep. Guillou et al. had cells in extensional flow region for < 5 ms (70 µm wide, 30 µm deep) and likely didn’t achieve quasi-steady cell strain, though cell strain vs. time for single cells was not reported, rather average maximum cell strain with flow rate [8]. Assuming the size of the constant extensional strain rate region maintains a 1/3 proportion to channel width, we can estimate the constant extensional strain rate region for these two studies was *r* ≤ 12 and 23 µm radius and therefore had only 1.2 – 4.6 times larger radius than the cells. This area was apparently too small and the fluid velocities too high for the cells with comparable shear moduli to agarose to reach a steady deformation state. Based on our study, the cells might require around 10 ms of dwell time in extensional flow to reach this state. In high Reynolds number experimental studies, 10 – 20 µm diameter cells in a 60-µm-wide, 30-µm-deep device dwelled only 20 – 30 µs in the central cross-slot region [19] and did not report a strain plateau. It is important to emphasize that these previous studies discriminated different cell groups or the mechanical effects of pharmacological agents on cell stiffness without achieving a quasi-steady cell strain. Furthermore, Cytovale offers an FDA-approved commercial product for early sepsis detection based on a high Reynolds number cross-slot device [21]. Quantitative biomechanical comparisons between groups using the cross-slot have been proven informative; but without achieving a quasi-steady plateau, measuring accurate and precise linear viscoelastic mechanical properties is more challenging.

Our large channel cross-slot implementation increased the centerline offset threshold below which particles achieve a strain plateau, thereby making quasi-steady particle deformation states during transient device operation more practical. Lu et al. simulated similar conditions to the previous cross-slot studies, studying the transient deformation of a capsule through a cross-slot with a square cross-section using a capsule diameter to channel width ratio of 0.4, e.g., a 15-µm diameter cell in a *w* = 37.5-µm square channel [22]. The authors predicted a centerline offset threshold of ≤ 0.01*w* in the straight inlet channel to achieve a quasi-steady strain state, which works out to less than 1 µm offset for the channel dimensions used in Refs. [5,8,19]. In experiments, this off-center tolerance seems too tight for the cross-slot channel dimensions used previously given that the data from the previous studies did not achieve a cell strain plateau. By extending the hyperbola fitted to particle trajectories upstream into the inlet channel, we calculated the offsets of the strain-plateaued particles in the straight inlet channels from the four flow rates to compare to the predictions of Lu et al. (**Figure S11, S16**). Most incoming particle offsets were ≤ 3 µm. Our choice of larger the device dimensions not only increased the area of uniform extensional flow region but also allowed for easier achievement of quasi-steady particle strains under experimental conditions.

Our study suggests design and particle analysis criteria guidelines for achieving a quasi-steady strain plateau for spherical particles that could yield more accurate mechanical property measurements. Firstly, the device channel width should be 6 or more times larger than the largest particle diameters. By conservatively selecting ranges where plateaued particles do not overlap with non-plateaued particles (**Figure 6**) the following analysis criteria could be applied to various biomaterials to select for objects that achieve a strain plateau: the object’s minimum distance to the stagnation point is less than or equal to the object’s diameter and the normalized change in velocity from inlet to stagnation point is ≤ 0.85. The following guidelines may depend more closely on the specific material and device dimensions: For 0.5% w/v agarose microparticles, particles that achieve offsets from the center streamline of ≤ 8 µm ∼ 0.03*w* when entering the constant extensional strain rate region and within this region dwelling ≥ 12.5 ms and traveling ≥ 145 µm = 0.5*w* would likely reach a quasi-steady strain state.

Our agarose microparticle elastic material parameter values were similar to a previous study, even with moderate strain variation within the particle strain plateaus due to low-contrast particles and further mechanical model validation ongoing. 0.5% w/w agarose microparticles were used to validate the real-time deformability cytometry technique’s measurement of particle elastic stiffness [47]. The reported agarose microparticles Young’s modulus was *E* = 2150 ± 1100 kPa, equivalent to shear modulus of *G* = 717 Pa assuming an incompressible material. Our measured range of shear elastic stiffness *G*_0_ = 1250 – 2250 Pa was somewhat higher but still reasonably close, thus encouraging further technique refinement. Future work to validate our approach could use microfluidic droplet-generated microparticles that are higher contrast to ensure more uniform particle material properties and improved edge detection accuracy. Our technique can also be used to explore particle relaxation after extensional flow as particles travel in the outlet channels by rotating the camera 90°. With the JetCam high-speed camera, flow rates, and cross-slot dimensions used here, only one measurement is possible, creep or relaxation, but not both, due to region-of-interest size limitations. Armistead et al. reported that cell elongation and relaxation strains had different exponential time constants [5], so the relaxation behavior seems important to measure as well.

The device dimensions, operating conditions, and strain plateau criteria should be tailored for each new material. Here, the proposed criteria are for 0.5% w/v agarose particles and similar soft biomaterials. Objects with different stiffness from soft agarose will require scaling of the characteristic hydrodynamic shear force *μ*Ω by tuning the fluid viscosity and flow rate. In laminar flow microfluidics, fluid velocity field quantities usually scale proportionally with flow rate, but PIV experiments are recommended for confirming flow conditions. Applying the design criteria suggested from this study on spherical particles to less spherical objects will likely require larger ‘factors of safety’ to achieve strain plateaus. Very spherical particles were compact and therefore less perturbed by outgoing flow at the unstable stagnation point location. We have found in preliminary experiments of ∼50- and ∼70-µm-equivalent-diameter cell clusters that were not perfect spheres first rigidly rotated and rearranged before stretching in extensional flow (same cross-slot device 290 µm wide, 265 µm deep, flow rate 4.5 mL/hr, Ω ∼ 144 s^-1^). Furthermore, the larger 70 µm clusters dwelled 20% less time in the cross-slot region compared to the 50 µm clusters because segments of the bigger non-symmetric shapes were more easily entrained in the outgoing fluid. Neither size cluster achieved a quasi-steady strain state, though another factor could be that the flow rates were too high for clusters that seem to be relatively softer than single cells [48]. Cell cluster microfluidic creep tests would likely be more successful if the channel width was made at least 500 µm wide, giving a channel width to particle effective diameter discrepancy of 7, or even up to 700 µm to reach a 700:70 = 10 ratio.

The distance from the cross-slot’s stagnation point is an important metric for assessing particle measurement quality. While the stagnation point location might be determined from the channel geometry alone, channel imperfections and system pressure fluctuations can perturb the fluid flow and make the stagnation point location known with less certainty. Using the equations of planar extensional flow, which was very well behaved within 100 µm of the stagnation point (**Figure 2, S3, S4**), the effective stagnation point for each particle can be assessed from a hyperbolic fit to the particle’s trajectory points. A good hyperbolic fit to the particle trajectory could serve as an additional quality check that particles are experiencing the desired extensional flow and therefore closely adhering to the mechanical model’s assumptions, thereby giving more confidence to the particle mechanical property measurements.

Our cross-slot implementation’s advantages over other microfluidic techniques that measure mechanical properties of microscale spherical objects arise from its large channel dimensions and stagnation point feature. Our technique operated in the fluid Stokes flow regime (low Reynolds number, negligible fluid inertia), particle linear viscoelasticity regime (small particle strain), and put the walls relatively far away (limited wall effects). These choices allowed for a proposed mechanical model that was a single analytical equation for determining particle viscoelastic material properties from the observed strain time history. Therefore, the data analysis of these experiments was simpler and more accessible to other research groups than those with close channel walls that require mechanical models implemented in numerical simulations. Previous cross-slot studies have demonstrated that it is challenging in practice to sufficiently center cells so that they reach a quasi-steady strain state in the extensional flow region near the stagnation point. The wide channels in our device design created a larger constant extensional strain rate region where more particles were likely to dwell long enough to reach a quasi-steady deformation state. With flow focusing, we observed that 50% of particles that were in-focus vertically were sufficiently centered laterally to closely approach the stagnation point and achieve a strain plateau. This yield could be improved with upstream device engineering that focuses more particles to the device cross-section geometric center. Relatively large channel dimensions are also less likely to lead to clogging when handling heterogenous mixtures of cell populations. The presence of the stagnation point means particle speeds were low when being stretched the most, which meant less blurring and the potential for more accurate shape measurements.

The large cross-slot channel dimensions have drawbacks related to measurement throughput and yield. Compared to finite Reynolds number techniques, low Reynolds number techniques like the one in this work will have lower measurement throughputs by 1 – 3 orders of magnitude. We measured 1 – 2 strain-plateaued particles per second that were good candidates for viscoelastic mechanical property measurements. The low Reynolds number real-time deformability cytometry technique measures single cells elastic or viscoelastic properties at throughputs around 100 cells per second [27–29]. The finite Reynolds number cross-slot-based deformability cytometry technique has mechanophenotyped single cells based on relative deformability at measurement rates of over 1000 cells per second [3,19,20]. Presumably in those techniques with close channel walls, most cells were in-focus and suitable for shape measurement. In our technique, roughly 25% of particles were focused at the midchannel height and sufficiently centered to be a good candidate for mechanical property measurements, meaning 75% of the sample was not measurable. More upstream device engineering to focus particles in both channel width and channel height directions would improve yield and throughput. Finite Reynolds number devices have the advantage of inertial flow focusing to center objects vertically and horizontally [49,50], which will lead to high measurement yield along with high throughputs, though at the cost of more challenging mechanical modeling to measure mechanical properties. At low Reynolds number, a sequence of converging/diverging channels used in real-time deformability cytometry [51,52] seems to center particles well. In droplet studies, it has been observed that a channel width narrow-to-wide change better aligned droplets at the center compared to a wide-to-narrow channel width transition [46]. Similar design choices could be implemented to improve our technique. Despite the relatively low throughput and yield for microfluidic techniques, our cross-slot microfluidic creep test is much faster than traditional micromechanical techniques like atomic force microscopy and micropipette aspiration, which have measurement throughputs of approximately 1 cell per minute [53].

Our cross-slot implementation shared analysis similarities with unidirectional techniques in which extensional flow was generated from changing channel width to study droplet interfacial tension [45,46,54] and hyperbolic channel walls to study cell mechanics [29,55–58]. In our approach, particles entered and left the frame in perpendicular directions while in these experiments, particles traveled in the same direction. In both unidirectional and perpendicular trajectory cases, the deforming hydrodynamic stresses acting on the particle, cell, or droplet depended on the object’s position in the device. It can be more convenient to have the analysis perspective that position within the device is the independent variable and other quantities (strain/deformation, elapsed time) are dependent on position. In the unidirectional devices, cells and droplets traveled at high speeds. The opposing flow configuration in this work had the advantage of low fluid and particle velocities around the stagnation point that facilitated easier particle shape measurements.

## CONCLUSIONS

High-throughput mechanical property measurements of biological objects like cells and cell clusters is important for practical use of these technologies in a clinical setting. Accuracy of mechanical property measurements and agreement between values reported by research groups has been a challenge. The cross-slot microfluidic device without active particle trapping could be a good compromise between accuracy and high throughput that is relatively easy to implement for many research groups. Here we demonstrated experimental conditions in which agarose microparticles of similar size to cells achieved the quasi-steady deformation state as they dwelled near the stagnation point. This transient strain plateau accurately captures the elastic response of spherical particles according to simulation predictions [22], thereby facilitating both accurate and rapid measurements. This work provides researchers with cross-slot design recommendations and particle strain plateau criteria for performing linear viscoelastic mechanical property measurements of single quasi-spherical objects. The proposed mechanical model is a starting point, and more validation work is needed. By providing a cross-slot technique that balances ease-of-use and potential accuracy, we aim to contribute to the micromechanical measurement field moving towards more standardized and repeatable measurement protocols and ultimately the clinical translation of biomechanics and mechanobiology knowledge to improve human health.

## Supporting information

Supporting Information doc

## Notes

### Competing Interest Statement

The authors have declared no competing interest.

