## Supporting Information doc for "Microfluidic creep experiment for measuring linear viscoelastic mechanical properties of microparticles in a cross-slot extensional flow device"

The supporting information includes:

**Viscosity of Aqueous Methyl Cellulose**

**Figure S1** Aqueous methyl cellulose viscosity.

**Table S1** Low-shear viscosity measurements.

**Particle Equilibrium Shape in No-Flow Conditions**

**Figure S2** Particle equilibrium shape in no-flow conditions.

**Particle Image Velocimetry**

**Constant Extensional Strain Rate Region**

**Figure S3** PIV central cross-slot region analysis for determining the constant extensional strain rate region.

**Figure S4** Velocity magnitude analysis results in 10-μm-wide bins in central cross-slot region.

**Figure S5** Constant extensional strain rate region analysis at 40x magnification

**Where Extensional Flow Begins**

**Figure S6** Velocity gradient components from the PIV data row nearest the centerline (on the *x* axis) and near-centerline data rows from a representative 100 μL/hr PIV experiments.

**Comparison of Aqueous Methyl Cellulose to a Newtonian Fluid**

**Figure S7** Comparison of straight channel Poiseuille flow profile for the shear-thinning 0.85% w/v aqueous methyl cellulose and Newtonian 0.394 M aqueous sucrose.

**Table S2** Summary of particle image velocimetry data from the 290-μm-wide, 265-μm-deep cross-slot device used in the hydrogel microfluidic creep experiments.

##### Deformed Particle Shape in Straight Channel

**Figure S8** Analytical predictions of particle deformation in Poiseuille flow [3] for a representative 50-μm particle from the experiments.

##### Particle Mechanical Model

**Figure S9** Fitted Kelvin Voigt parameters for strain-plateauing and non-plateauing particles.

**Figure S10** Particle inlet speed and average strain in inlet for strain-plateauing and non-plateauing particles.

**Figure S11** Particle offsets in straight inlet channel for strain-plateauing and non-plateauing particles.

##### Particles with Plateaus

**Figure S12** Particle trajectories and strain for flow rate 6.5 mL/hr, $N$ = 21.

**Figure S13** Particle trajectories and strain for flow rate 7.5 mL/hr, $N$ = 40.

**Figure S14** Particle trajectories and strain for flow rate 8.5 mL/hr, $N$ = 37.

**Figure S15** Particle trajectories and strain for flow rate 9.5 mL/hr, $N$ = 42.

**Figure S16** Particle offsets in straight inlet channel for four flow rates.

### Viscosity of Aqueous Methyl Cellulose

The viscosity of the 0.85% w/v methyl cellulose (Thermo Scientific Chemicals, CAS 9004-67-5, molecular weight 454.513 g/mol) solution in which the agarose hydrogel particles were suspended was an input to the mechanical model that extracted particle viscoelastic material properties from the microfluidic creep experiments. Viscosity was measured on a TA Instruments Discovery Hybrid series rheometer with a parallel plage geometry (40 mm diameter). Viscosity was determined from a shear rate sweep protocol between 1 – 100 or 1000 $s^{-1}$ strain rates with 10 pointes per decade. Three technical replicate were performed for each fluid case.

All the aqueous methyl cellulose used in the hydrogel microfluidic creep experiments performed on two separate days was prepared from a single batch (Batch 1). Most of Batch 1 was filtered through a 0.2-μm-pore PVDF filter. A portion (~1/3) of the fluid used in the first experiment day was unfiltered Batch fluid. These two versions of Batch 1 were measured for viscosity over the shear rate range 1 – 100 $s^{-1}$. A second batch of 0.85% w/v aqueous methyl cellulose was prepared and measured over a larger shear rate range 1 – 1000 $s^{-1}$ to confirm the power law behavior at high shear rates observed previously [1]. This Batch 2 was vacuum filtered using Whatman Qualitative Filter Paper, Grade 5 (pore size ~2.5 μm).

Our results show that methyl cellulose was shear thinning (viscosity decreases with increasing shear rate), as reported by others [1]. Within the precision of the instrument and the protocol, the viscosity for the three fluid cases (0.2-μm-PVDF filtered, unfiltered, cellulose filter paper) did not differ significantly. The fluid sample filtered with the smallest 0.2-μm pore size had the least variability between the three technical replicates compared to the unfiltered and larger pore size filtered cases. This is consistent with the assumption that a smaller pore size filter will yield a more homogenous fluid.

The viscosity used in the mechanical model (Equation (6) in manuscript) was taken to be the average viscosity of Batch 1 filtered and unfiltered at shear rates 10 $s^{-1}$ and 12.6 $s^{-1}$, an estimate the fluid's zero-shear viscosity. We assumed low shear rates (off-diagonal velocity gradient terms $\partial u/\partial y = \partial v/\partial x \sim 0$) in the extensional flow region based on the PIV measurements (**Figure S6**), so the effective fluid viscosity felt by the deforming particles would be close the fluid's zero-shear viscosity. A previous study of aqueous methyl cellulose shear rheology [1] showed that parallel plate geometry viscosity measurements around 10 $s^{-1}$, the lowest shear rate where this geometry is accurate, agree closely with the concentric cylinder measurements at low shear rates that showed aqueous methyl cellulose reaches a viscosity plateau below 10 $s^{-1}$.

Our final estimate of the effective viscosity was $\mu$ ~ 110 mPa·s. This was consistent with the zero-shear viscosity of 88 mPa·s reported for slightly less concentrated 0.83% w/v methyl cellulose in phosphate buffered saline [1] given methyl cellulose's rapid increase of viscosity with polymer concentration (see methyl cellulose product information).

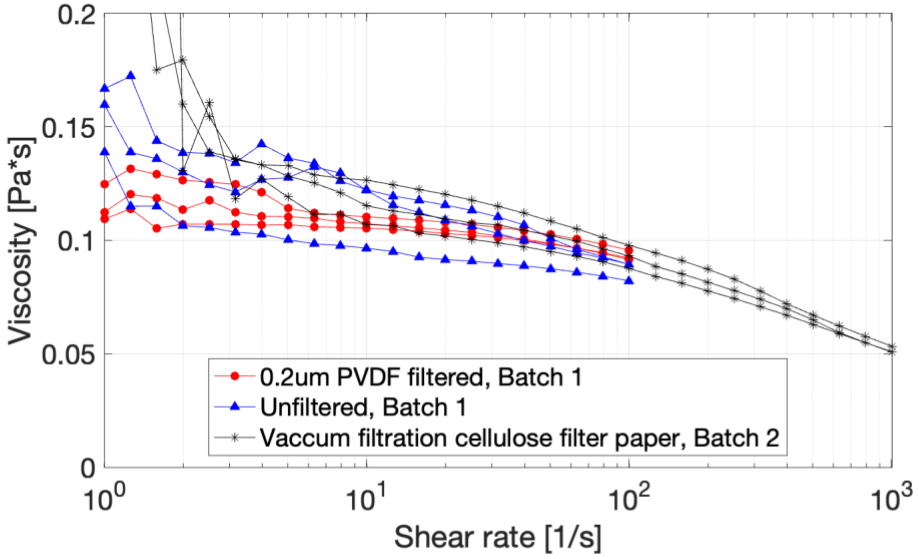


**Figure S1  Aqueous methyl cellulose viscosity.** Viscosity of 0.85% w/v aqueous methyl cellulose with shear rate with and without filtering. Three technical replicates were performed for each filtration case. The red and blue curves show the viscosity data of the fluid used in the hydrogel experiments that took place over two days.

| | | Shear Rate [1/s] | |
|---|---|---|---|
| Viscosity [Pa·s] | Technical Replicate | 9.99985 | 12.5893 |
| 0.2-µm-pore Filtered | 1 | 0.107249 | 0.106392 |
| | 2 | 0.105333 | 0.104739 |
| | 3 | 0.110371 | 0.109625 |
| Unfiltered | 1 | 0.122249 | 0.119509 |
| | 2 | 0.121974 | 0.115474 |
| | 3 | 0.0964929 | 0.0950883 |

| | |
|---|---|
| Average Viscosity [Pa·s] | 0.110 |
| Standard Deviation [Pa·s] | 0.009 |

**Table S1. Low-shear viscosity measurements** of Batch 1 0.85% (w/w) aqueous methyl cellulose that were averaged to estimate the zero-shear viscosity of this fluid.

### Particle Equilibrium Shape in No-Flow Conditions

Particles were imaged in a still fluid over four seconds at 50 frames per second or 200 frames per particle. Particle shape was assessed in the same way as the microfluidic experiments: The contour was detected by finding the maximum intensity gradient along rays drawn outwards from the center, and then an ellipse was fit to the contour. The strain was defined as $\varepsilon = (b - a)/(b + a)$ where $b$ is the ellipse semi-axis in the vertical direction and $a$ is the ellipse axis in the horizontal direction. Our final estimate of the no-flow particle equilibrium strain as measured with our microscopy system and edge detection algorithm is 0.006 ± 0.003 (mean ± standard deviation, *N* = 36).

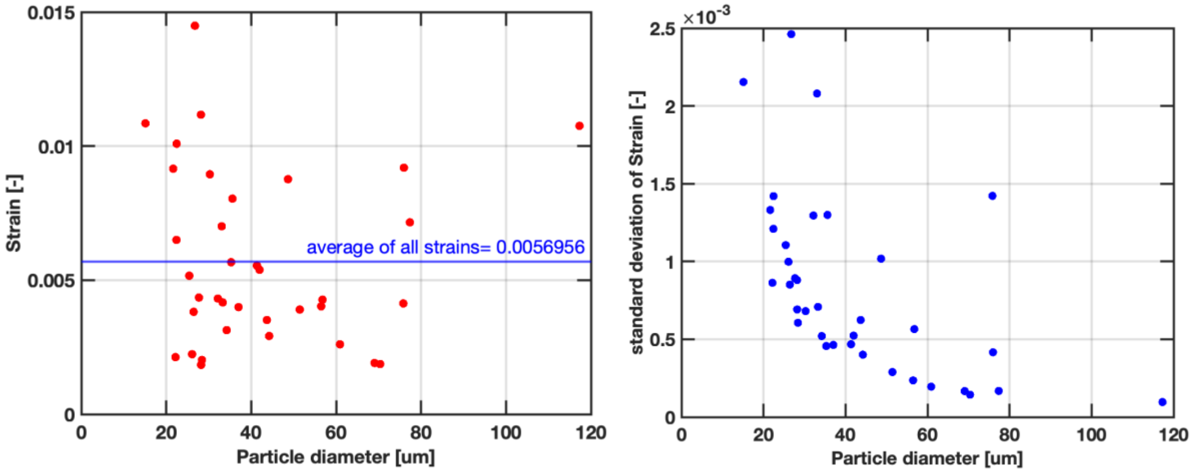


**Figure S2 Particle equilibrium shape in no-flow conditions.** Left: The average strain over 200 frames for *N* = 36 particles. The grand average strain is 0.006 with a standard deviation of 0.003. Right: The standard deviation of the individual particle strains over 200 frame movies.

### Particle Image Velocimetry

#### Constant Extensional Strain Rate Region

The radius of the constant extensional flow region centered at the stagnation point was taken to be the region where the standard deviation of the velocity magnitude within 10-µm-wide bins did not vary by more than 10% of the average standard deviation for bins in the region $0 \leq r \leq 50$ µm. $r$ is the distance from the stagnation point.

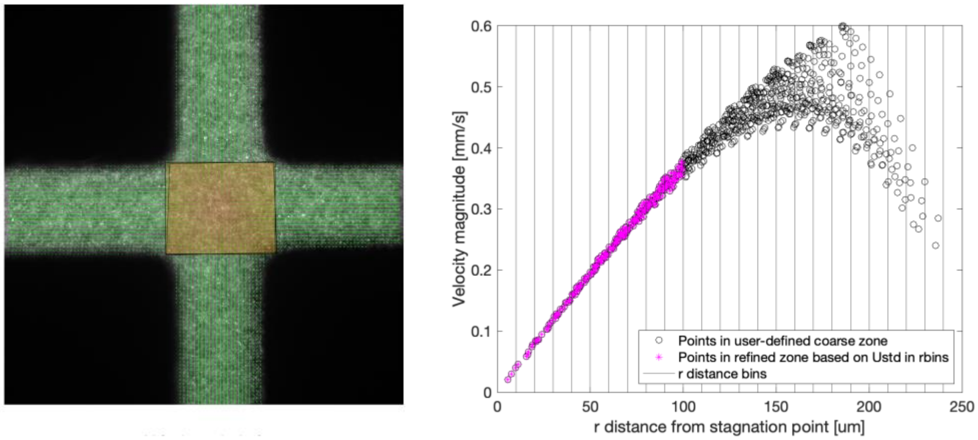


**Figure S3 PIV central cross-slot region analysis for determining the constant extensional strain rate region.** Left: The rectangle highlights the central cross-slot region where velocity vector magnitude $|\mathbf{v}|$ vs. distance from the stagnation point $r$ was analyzed to determine the constant extensional strain rate region. Right: $|\mathbf{v}|$ vs. $r$ scatter plot. The standard deviation, a measure of the width of scattered data points around the linear regression trend line, was very uniform for data points within $r \leq 100$ µm.

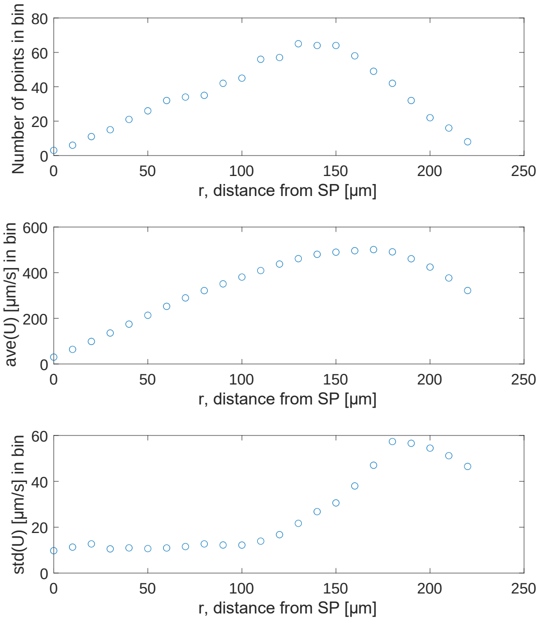


**Figure S4 Velocity magnitude analysis results in 10-µm-wide bins in central cross-slot region.** Top: Number of PIV velocity vectors within each bin. Middle: Average velocity vector magnitude within the bins. Bottom: Standard deviation of the velocity vector magnitude within the bins. There was an increase in standard deviation starting at 100 µm from the stagnation point, marking the edge of the constant extensional strain rate region.

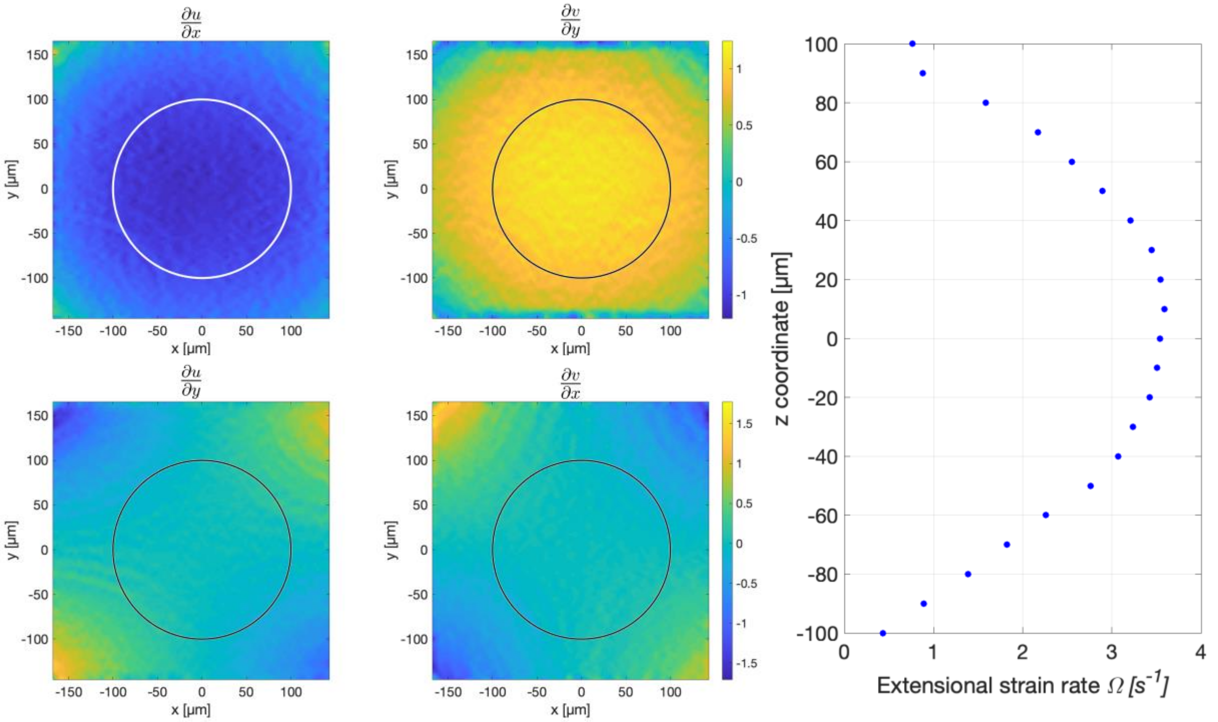


**Figure S5 Constant extensional strain rate region analysis at 40x magnification.** Left: Velocity gradient components. Circle show the constant extensional strain rate circular region with radius 100 μm centered at the stagnation point. Right: Variation of the constant extensional strain rate $\Omega$ with *z*, the coordinate direction in the direction of gravity. The constant strain rate $\Omega$ is the slope of the linear regression fit of velocity magnitude against distance from the stagnation point for PIV data points within 100 μm from the stagnation point.

#### Where Extensional Flow Begins

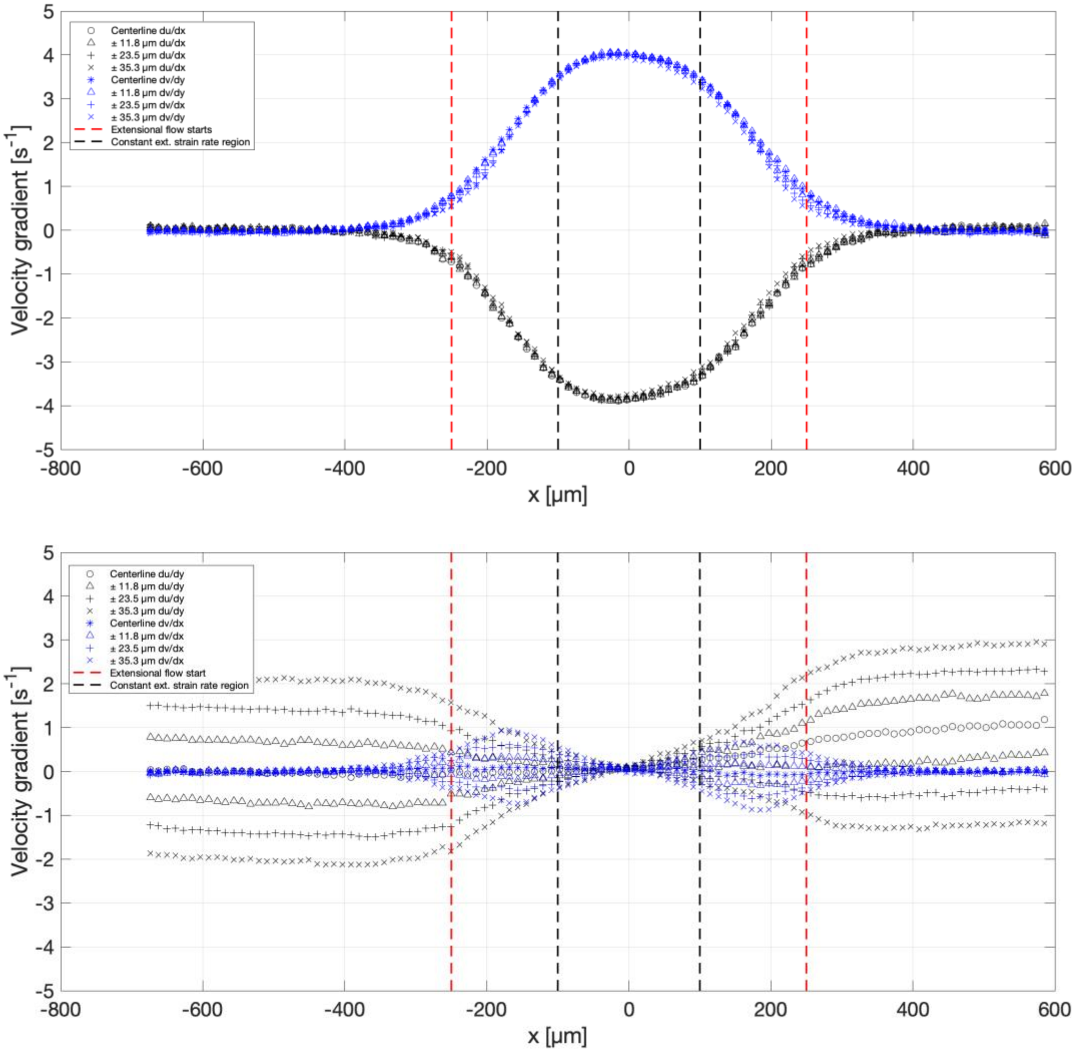


**Figure S6 Velocity gradient components from the PIV data row nearest the centerline (on the *x* axis) and near-centerline data rows from a representative 100 µL/hr PIV experiments.** Top: Diagonal velocity gradient components $\partial u/\partial x$ and $\partial v/\partial y$. Bottom: Off-diagonal velocity gradient components $\partial u/\partial y$ and $\partial v/\partial x$. Vertical red dashed lines mark 250 µm from the stagnation point where the diagonal and off-diagonal velocity gradient component have appreciably changed from their values in the straight channels. The vertical black dashed lines bracket the constant extensional strain rate region where $-\partial u/\partial x \sim \partial v/\partial y \sim \Omega$ and $\partial u/\partial y \sim \partial v/\partial x \sim 0$.

### Comparison of Aqueous Methyl Cellulose to a Newtonian Fluid

The degree to which shear thinning methyl cellulose departed from Newtonian behavior was assessed by comparing the parabolic flow profiles in the straight channels for PIV measurements using methyl cellulose to 1 μm particles suspended in sucrose, which is basically water and a Newtonian fluid. A suspension of 0.005% v/v 1-μm fluorescent polystyrene spheres in 0.394 M aqueous sucrose (Sigma) was infused in the microfluidic device. The parabolic flow profiles $u(y)$ and $v(x)$ in the straight inlet and outlet channels were compared between the two fluids at flow rates of 75 and 100 μL/hr.

A shear thinning fluid is expected to have a blunter parabolic profile near the device middle and higher gradients near the wall. The bluntness of the parabolic profiles was compared using the focal lengths $1/a$ of the fitted parabolas $u(y) = ay^2 + by + c$ in the straight channels. A shorter focal length indicated a blunter parabola. The flow profiles were very similar between the shear thinning methyl cellulose and the Newtonian aqueous sucrose with a lot of overlap in the data (**Figure S7**). Looking at averages, the average focal length is shorter for the aqueous methyl cellulose compared to the Newtonian aqueous sucrose, but only by a small amount. Thus, the shear-thinning aqueous methyl cellulose was slightly blunter than the Newtonian aqueous sucrose but not by a significant amount. The resolution of the PIV measurements was too low to assess the fluid fields near the channel walls.

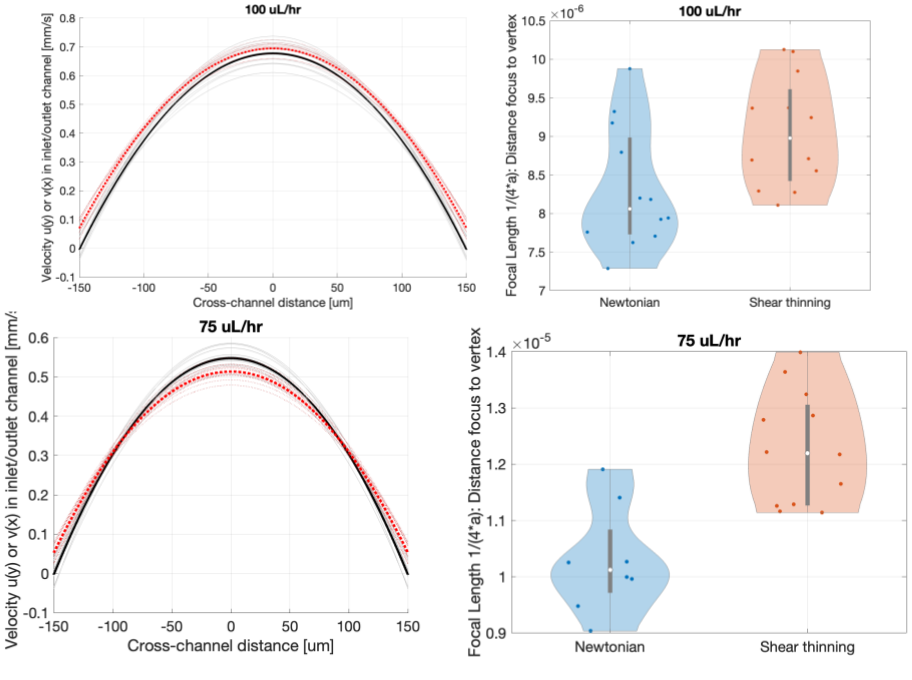


**Figure S7 Comparison of straight channel Poiseuille flow profile for the shear-thinning 0.85% w/v aqueous methyl cellulose and Newtonian 0.394 M aqueous sucrose.** Top: 100 μL/hr experiments. Bottom: 75 μL/hr experiments. Red line: average parabola fit to methyl cellulose experiments. Black line: Average parabola fit to aqueous sucrose Transparent black and red lines: Parabola fits to individual straight channels for both fluids.

**Table S2 Summary of particle image velocimetry data from the 290-µm-wide, 265-µm-deep cross-slot device used in the hydrogel microfluidic creep experiments.** Three technical replicates were conducted per flow rate with 10x magnification. The radius of the constant extensional flow region centered at the stagnation point was taken to be the region where the standard deviation of the velocity magnitude within 10-µm-wide bins did not vary by more than 10% of the average standard deviation for bins in the region $0 \le r \le 50$ µm. $r$ is the distance from the stagnation point. The theoretical maximum speed of the Poiseuille flow in the straight, rectangular cross-section channels was computed from the infinite Fourier series solution presented truncated to 16 terms [2].

| Flow rate in each inlet channel [µL/hr] | PIV data min(\|v\|) [µm/s] | Average std(\|v\|) in 10-µm-wide bins from $0 \le r \le 50$µm [µm/s] | Radius of constant extensional strain rate $\Omega$ [µm] | Constant extensional strain rate $\Omega = d\|v\|/dr$ over points $r \le 100$µm [1/s] | Average constant extensional strain rate $\Omega$ (N = 3) [1/s] Figure 1F | Standard deviation constant extensional strain rate $\Omega$ ($N$ = 3) [1/s] | Maximum Poiseuille flow speed $u(y)$, $v(x)$ parabola fits, averaged 4 straight channels [mm/s] | Maximum Poiseuille flow speed, averaged ($N$ = 3) [mm/s] Figure 1H | Theoretical maximum Poiseuille flow speed [mm/s] | Max inlet speed: % difference theory to PIV experiment |
|---|---|---|---|---|---|---|---|---|---|---|
| 50 | 9.4 | 7.57 | 100 | 1.827 | | | 0.3566 | | | |
| 50 | 11.0 | 7.24 | 110 | 1.724 | **1.77** | 0.05 | 0.3432 | **0.3488** | 0.3785 | -7.8% |
| 50 | 6.9 | 6.77 | 100 | 1.771 | | | 0.3466 | | | |
| 75 | 17.3 | 22.33 | 190 | 2.586 | | | 0.5131 | | | |
| 75 | 10.4 | 15.39 | 120 | 2.578 | **2.60** | 0.03 | 0.5039 | **0.5154** | 0.5678 | -9.2% |
| 75 | 19.5 | 24.2 | 150 | 2.634 | | | 0.5293 | | | |
| 100 | 18.2 | 10.85 | 60 | 3.558 | | | 0.6910 | | | |
| 100 | 20.5 | 11.05 | 70 | 3.628 | **3.58** | 0.04 | 0.7054 | **0.6958** | 0.757 | -8.1% |
| 100 | 19.3 | 10.78 | 60 | 3.555 | | | 0.6911 | | | |
| 125 | 27.2 | 12.2 | 50 | 4.348 | | | 0.8473 | | | |
| 125 | 24.0 | 13.75 | 100 | 4.494 | **4.41** | 0.07 | 0.8748 | **0.8599** | 0.9463 | -9.1% |
| 125 | 23.0 | 13.03 | 100 | 4.399 | | | 0.8575 | | | |
| 150 | 29.8 | 19.08 | 100 | 5.404 | | | 1.0542 | | | |
| 150 | 35.9 | 16.52 | 70 | 5.416 | **5.38** | 0.05 | 1.0563 | **1.0501** | 1.1355 | -7.5% |
| 150 | 33.9 | 14.63 | 50 | 5.319 | | | 1.0398 | | | |

#### Deformed Particle Shape in Straight Channel

Before experiencing extensional flow in the central region of the cross-slot device, particles flowed through straight channel with a rectangular cross-section (here width ~290 μm, height ~ 265 μm). Thus, particles initially experienced pressure driven, unidirectional Poiseuille flow in the cross-slot device. Particles in Poiseuille flow will deform into a bullet or parachute shape that is elongated at the leading edge and flattened at the trailing edge [3]. Our particle edge detection algorithm detected a small amount of deformation of 0.5% w/v agarose microparticle, and here we checked if the Poiseuille flow was strong enough in our experiments to deform the particle towards a bullet shape.

Murata used perturbation theory to predict the deformation of an elastic solid sphere in unbounded linear flow fields [4]**.** Villone et al. interpreted Murata's result for elastic particles deforming in Poiseuille flow in circular cross-section channel [3,5], which for a particle centered on the channel's symmetry axis (central streamline) simplifies to:

$$r(\theta,\phi) = R_p\left(1 + 4\frac{\mu U}{2R_t G}\left[\frac{7}{16}\frac{R_p}{R_t}(5\cos^3\theta - 3\cos\theta)\right]\right) \qquad \text{(S1)}$$

Where $r(\theta,\phi)$ is a point on the surface of the deformed sphere in spherical coordinates (polar coordinate $0 \leq \theta \leq \pi$, azimuthal coordinate $0 \leq \phi \leq 2\pi$, flow direction and channel axis is $\theta = 0$), $R_p$ is the equilibrium radius of the elastic sphere under no forces, $\mu$ is the viscosity of the Newtonian fluid, $U = \frac{Q}{\pi R_t^2}$ is the average fluid velocity based on the infused flow rate $Q$ and cross-sectional area of the channel with radius $R_t$, and the sphere's shear elastic modulus $G$. Villone et al. found that their numerical simulations agreed well with Murata's theoretical predictions of deformed sphere shape for low confinement (the ratio $R_p/R_t$ is small) and weaker viscous forces (small capillary number for this scenario $Ca = \mu U/2R_t G$), as expected given that Murata's analysis assumes small deformation $(r(\theta,\phi) - R_p)/R_p \ll 1$ [5].

To use Equation 1 for a cylindrical channel geometry to estimate the deformation of particles in our experimental device with a rectangular cross section, we find an equivalent channel diameter for our experimental set up. The equivalent diameter yields the same pressure drop along the straight inlet channel for the same flow rate in the rectangular. Using the analytical expressions for flow rates in a pressure driven flow in cylindrical and rectangular geometries [2] and setting the pressure drop to be equal, we find the equivalent channel radius for the experimental rectangular channel to be:

$$R_{t,equiv} = \left(\frac{8}{\pi}\frac{h^3 w}{12}\left[1 - 6\frac{h}{w}\sum_{i=0}^{n}\lambda_n^{-5}\tanh\left(\frac{\lambda_n w}{h}\right)\right]\right)^{\frac{1}{4}} \qquad \text{(S2)}$$

Where *h* and *w* are the rectangular channel's height and width and $\lambda_n = (2n+1)\pi/2$. For *h* = 265 μm and *w* = 290 μm, $R_{t,\,equiv}$ ~ 151. 48 μm.

The expected deformation of a 50-μm diameter agarose particle in the straight inlet channel was evaluated from Equation S1 using parameters from the experiments (**Figure S7**). The particle's confinement ratio is $R_p/R_{t,equiv}$ = 25/151.48 ~ 0.165. For a flow rate of Q = 8.5 mL/hr, the average velocity in the equivalent circular cross section channel is $U = Q/\left(\pi R_{t,equiv}^2\right)$ ~ 0.033 m/s. The fluid's viscosity is approximately $\mu$ ~ 110 mPa*s, and the particle's shear elastic modulus is a typical value of $G_0$ = 1750 Pa. The elastic capillary number is small $Ca = \mu U/(2R_t\,G) \sim 0.007$, meaning that the fluid viscous forces were not very strong compared to the particle's stiffness that is resisting deformation by those viscous forces. An ellipse was fitted to the contour points spaced by 1 degree, the experimental contour point density. The fitted ellipse long and short axes were 50.0030 and 50.0014 μm, respectively. This resulted in a strain $\varepsilon$ = (50.0030 – 50.0014)/(50.0030 + 50.0014) = 1.65e-5, which was well below the detection limit of the particle edge detection algorithm.

Therefore, theoretical predictions suggested that the particles in low confinement in the experiments (confinement ratio typically ≤ 0.165) were not significantly deformed by the pressure-driven Poiseuille flow when well-centered. This means that the non-zero strain measured for particles upstream of the extensional flow was probably due to artifacts such as blurring due to fast movement and relatively slow camera exposure times and/or edge detection inaccuracies due to low signal-to-noise images rather than deformation due to the viscous fluid forces.

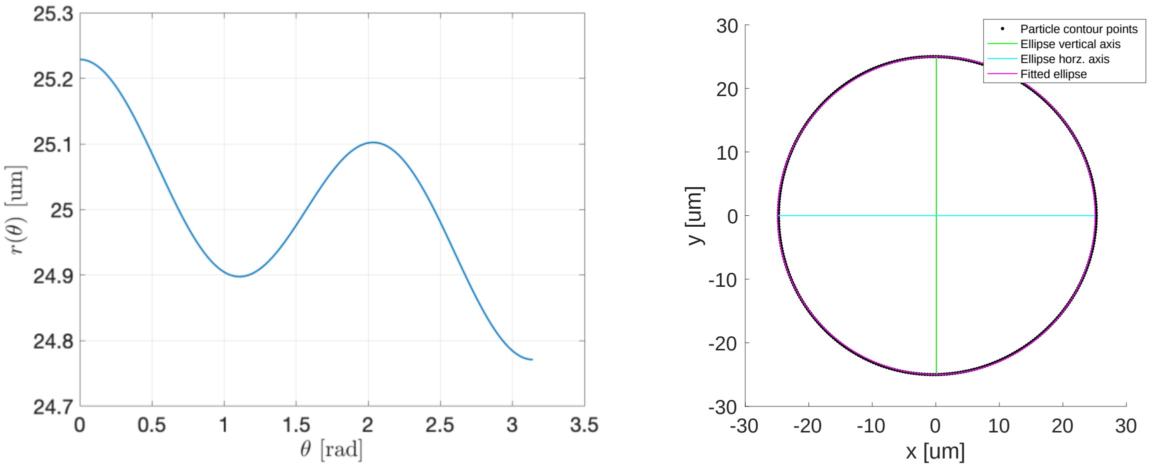


**Figure S8 Analytical predictions of particle deformation in Poiseuille flow [3] for a representative 50-μm particle from the experiments.** Left: Direct prediction of particle radius from Equation S1. Right: Predicted particle cross-section with fitted ellipse overlay.

### Particle Mechanical Model

Using the viscoelastic correspondence principle [6], the Laplace-transformed viscoelastic solution of a linear viscoelastic sphere deforming in planar extensional flow was obtained directly from the corresponding elastic solution. Murata solved the linear elastic, linear flow field problems for elastic sphere in an arbitrary linear flow field using a perturbation approach where the small variable was displacement from the equilibrium/initially spherical shape. From Murata's solutions for an incompressible elastic sphere deforming in a linear flows of a Newtonian fluid at low Reynolds [4], we obtained at the strain of an elastic sphere at its equator when deforming at the stagnation point of planar extensional flow:

$$\varepsilon = \frac{5}{2}\frac{\mu\Omega}{G} \tag{S3}$$

$\Omega$ is the uniform velocity field extensional strain rate, $\mu$ is the fluid Newtonian viscosity, and $G$ is the sphere's shear elastic modulus. To arrive at the solution of the viscoelastic problem, the quantities in Equation S3 were reinterpreted as their Laplace transforms. The elastic field variable, strain $\varepsilon$, is interpreted as $\bar{\varepsilon}(s)$ where $s$ is the transform parameter. For the boundary conditions, viscous extensional fluid traction boundary conditions were suddenly turned on as the microparticles approached the cross-slot's extensional flow region, and this was represented by the Heaviside unit step function $H(t)$. So, the elastic problem's $\Omega$ is interpreted as $\Omega H(t)$ for the viscoelastic problem and was replaced by its Laplace transform $\Omega/s$. The viscoelastic response function—here the relaxation modulus $G(t)$—was replaced by the transform parameter multiplied by its Laplace transform, $s\bar{G}(s)$. Making these replacements into the elastic solution Equation S3, the Laplace transform of the viscoelastic problem was:

$$\bar{\varepsilon}(s) = \frac{5}{2}\mu\frac{\Omega}{s}\frac{1}{s\bar{G}(s)} \tag{S4}$$

The choice of the viscoelastic response function $\bar{G}(s)$ depends on the material's constitutive law. The result for a viscoelastic sphere is obtained upon taking the inverse Laplace transform of Equation S4.

### Strain Plateauing Particles vs. No Strain Plateau Particles

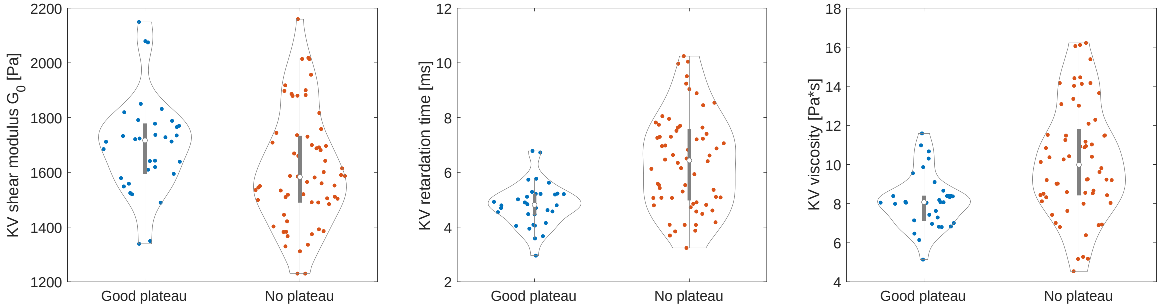


**Figure S9 Fitted Kelvin Voigt parameters for strain-plateauing and non-plateauing particles.** Comparison of the Kevlin Voigt mechanical properties for strain plateauing particles and non-plateauing particles. Left: Shear modulus $G_0$ was similar. Middle: Plateauing particles had a narrower range of retardation time $\tau$ than non-plateauing particles. Right: Platuing particles also had a tighter range of viscosity $\eta = G_0\tau$.

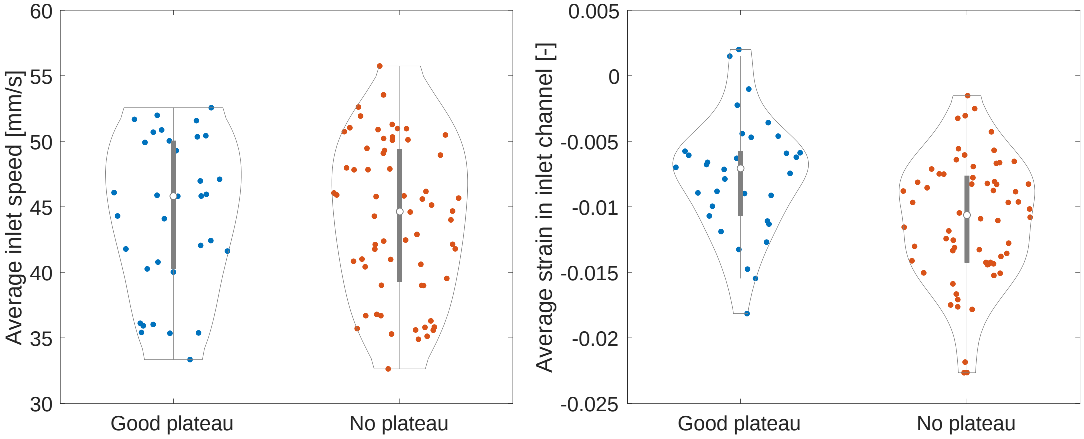


**Figure S10 Particle inlet speed and average strain in inlet for strain-plateauing and non-plateauing particles.** Left: Incoming particle speeds were similar. Right: Non-plateauing particles tended to have larger strains, but the accuracy of incoming strain should be improved before drawing further conclusions.

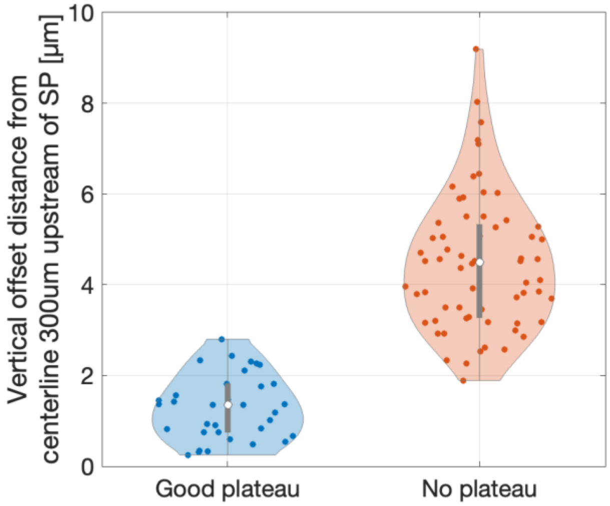


**Figure S11 Particle offsets in straight inlet channel for strain-plateauing and non-plateauing particles.** The particle offset from the center streamline in the inlet channel was estimated at the location 300 μm upstream of the stagnation point just before particles began to slow down and start turning on a hyperbolic streamline. The strain-plateauing particles had a tight range of offsets ≤ 3 μm lower than the non-plateauing particles ≥ 2 μm offsets.

### Particles with Plateaus

*In the following figures of particles that achieved a quasi-steady strain plateau:*
For all subfigures, the grayscale trajectory and strain marker colors correspond to the particle's offset from the center streamline upon entering the constant extensional strain rate region (black circle).
Top: Particle trajectories overlayed on a representative experimental movie frame. The particle data came from two different days for which the stagnation points to not perfectly align. The red dashed line marks 250 µm upstream of the stagnation point (from one of the days) where extensional flow starts.
Bottom Left and Right, the particle strain data were aligned by choosing *t* = 0 to correspond to the location 250 µm upstream of the stagnation point where the extensional flow starts (red dashed vertical line). The solid black vertical line marks when the particles entered the constant extensional strain region (*r* ≤ 100 µm), and the grayscale dashed vertical line marks where individual particles left this region.

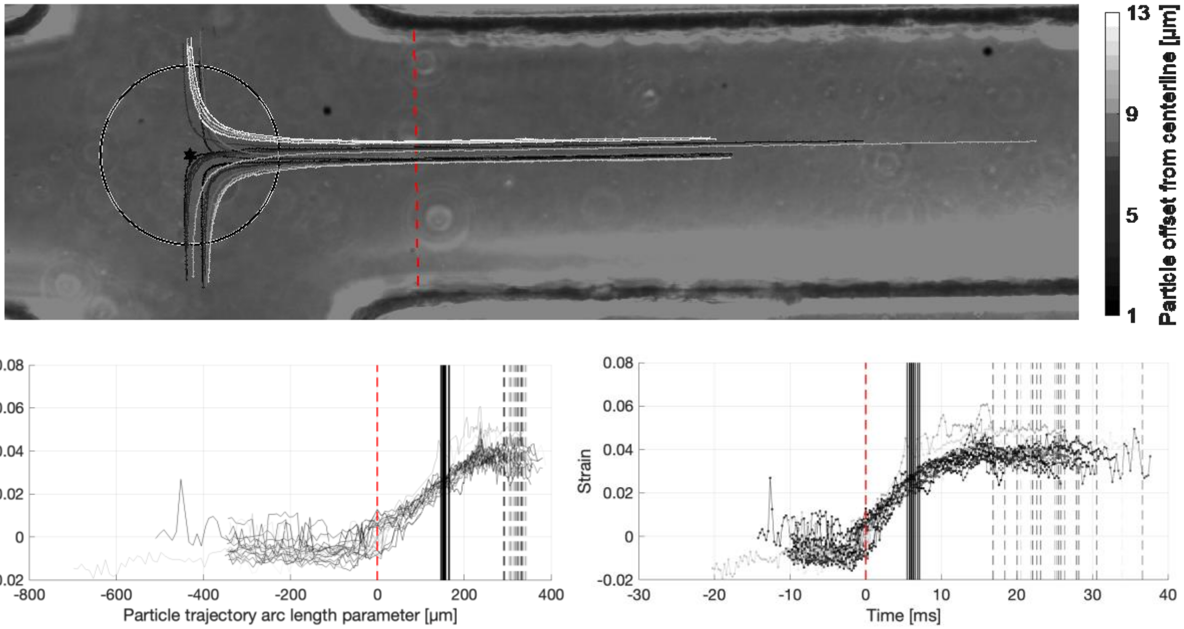


s

**Figure S12 Particle trajectories and strain for flow rate 6.5 mL/hr, *N* = 21.**

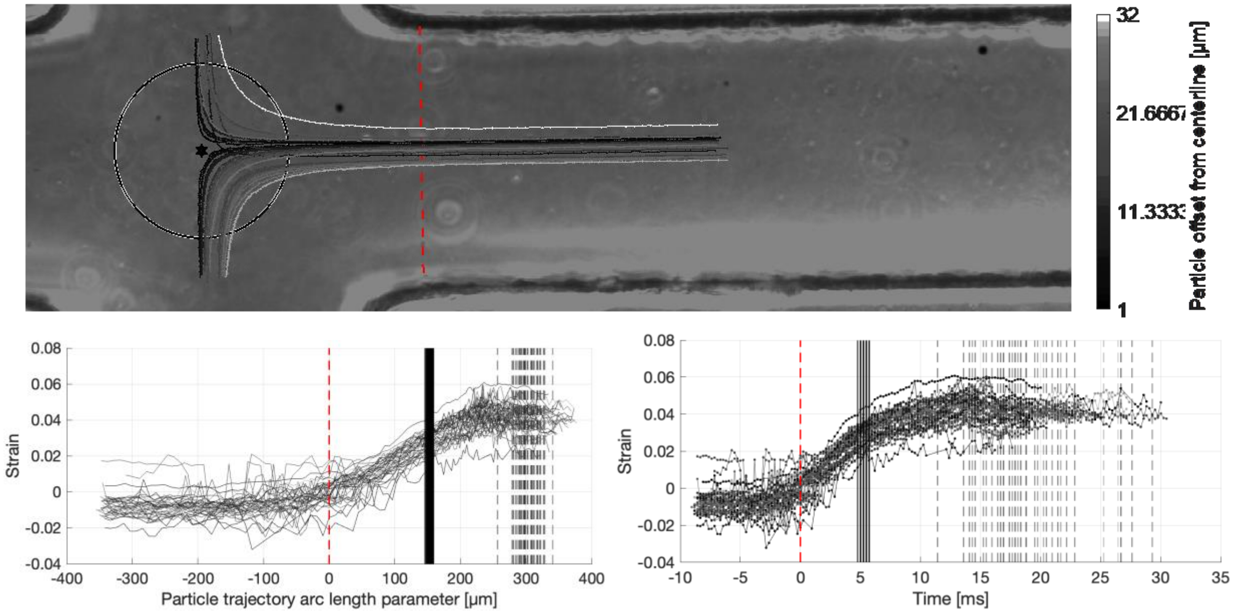


**Figure S13 Particle trajectories and strain for flow rate 7.5 mL/hr, *N* = 40.**

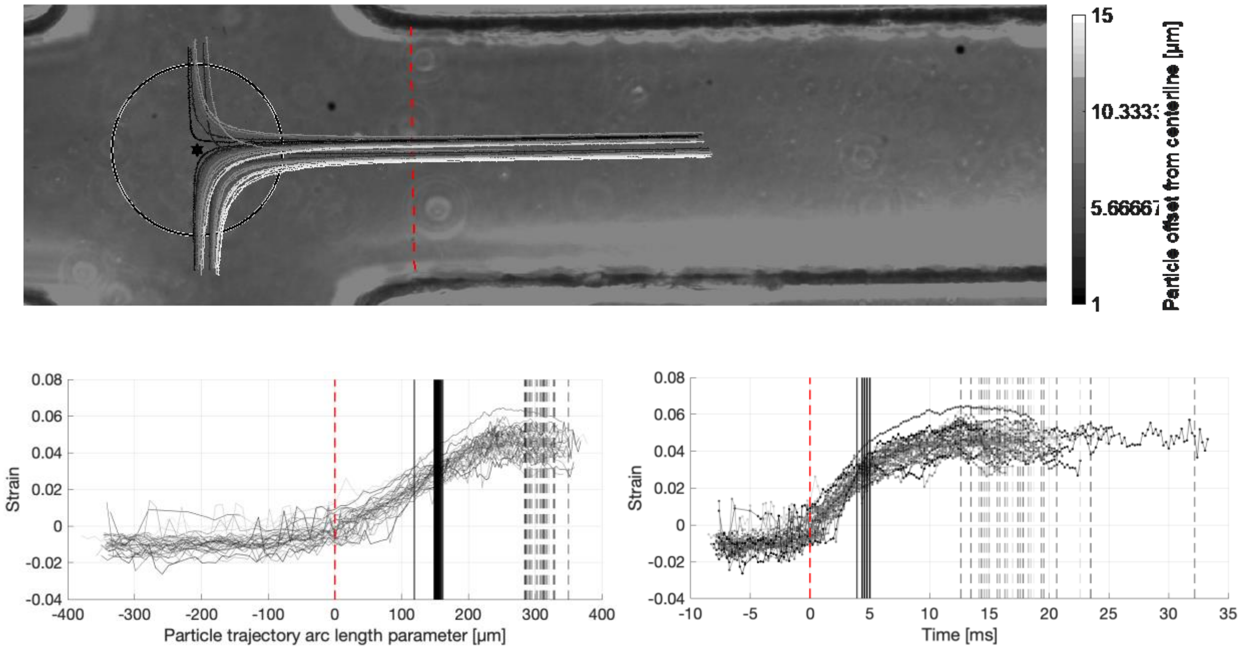


**Figure S14 Particle trajectories and strain for flow rate 8.5 mL/hr, *N* = 37.**

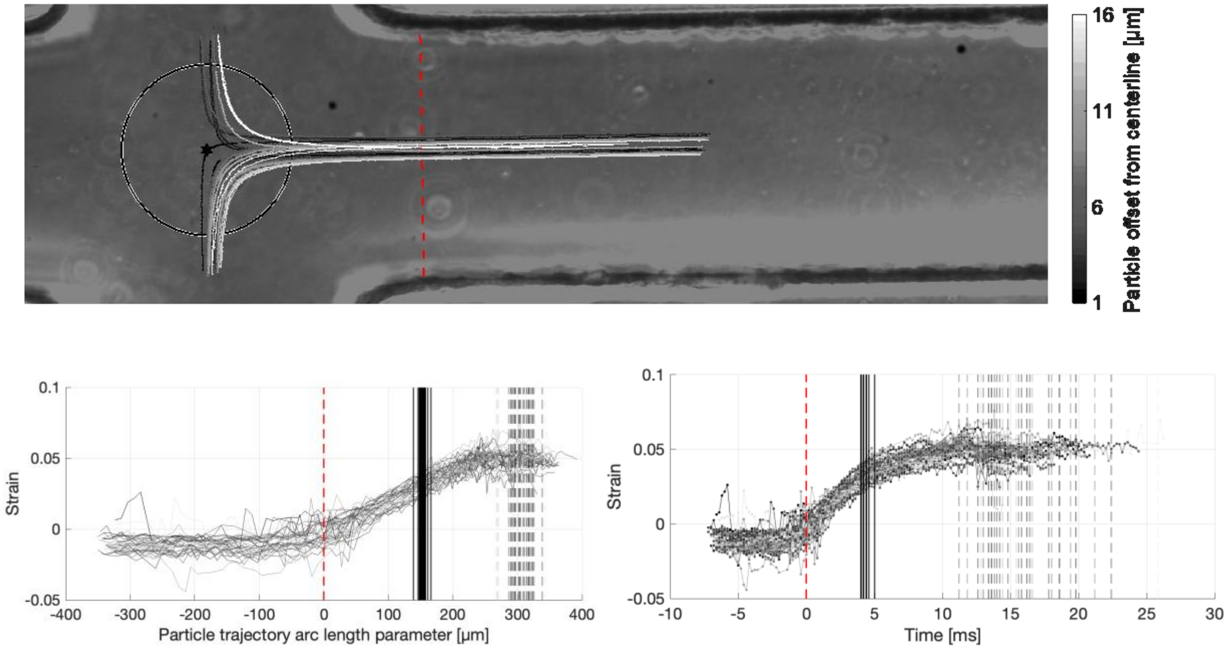


**Figure S15 Particle trajectories and strain for flow rate 9.5 mL/hr, *N* = 42.**

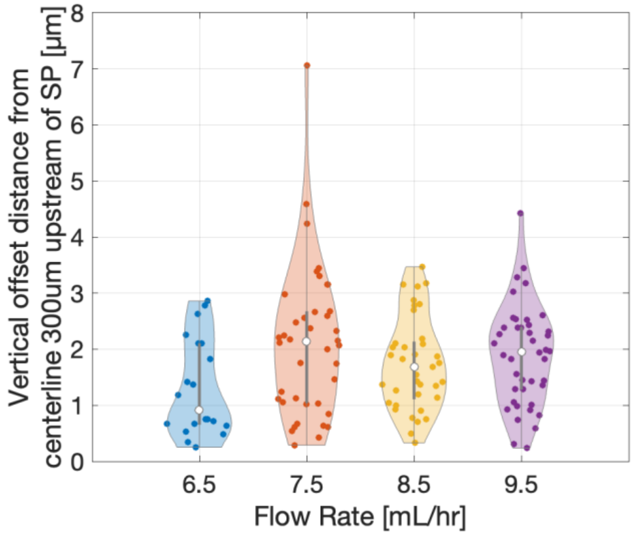


**Figure S16 Particle offsets in straight inlet channel for four flow rates.** The particle offset from the center streamline in the inlet channel was estimated at the location 300 µm upstream of the stagnation point just before particles began to slow down and start turning on a hyperbolic streamline. The strain-plateauing particles had a tight range of offsets ≤ 3 µm lower than the non-plateauing particles ≥ 2 µm offsets.
